# The proto-oncogene c-Src phosphorylates cGAS to reduce DNA binding and activation

**DOI:** 10.1101/2022.10.07.511364

**Authors:** William Dunker, Shivam A Zaver, Cameron J Howard, Joshua J Woodward

**Author notes:** These authors contributed equally to this work (William Dunker, Shivam A Zaver).

## Abstract

cGMP-AMP synthase (cGAS) is a DNA sensor and responsible for inducing an anti-tumor immune response. Recent studies reveal cGAS is frequently inhibited in cancer, and therapeutic targets to promote anti-tumor cGAS function remain elusive. c-Src is a proto-oncogene tyrosine kinase and is expressed at elevated levels in numerous cancers. Here, we demonstrate that c-Src expression in tumors from various cancers negatively correlates with innate immune gene expression and patient survival. We determine that c-Src restricts cGAS signaling in human cell lines through c-Src small molecule inhibitors, depletion, and overexpression. cGAS and c-Src interact in cells and *in vitro*, while c-Src directly inhibits cGAS enzymatic activity and DNA binding in a kinase-dependent manner. c-Src phosphorylates cGAS, and inhibition of cGAS Y248 phosphorylation partially reduces c-Src inhibition. Collectively, our study demonstrates that cGAS anti-tumor signaling is hindered by the proto-oncogene c-Src and describes how cancer-associated proteins can regulate the innate immune system.

## Introduction

The activation and regulation of the cell intrinsic innate immune system is pivotal for the induction of an anti-microbial response to an infection, for restricting endogenous molecules from promoting an auto-immune state, and for initiating cell death following catastrophic cellular damage. Germline-encoded pattern recognition receptors (PRRs) are frontline defense sensors that recognize distinct microbial pathogen-associated molecular patterns (PAMPs) and damage-associated molecular patterns (DAMPs) to initiate an immune response ^1^. One PRR class includes nucleic acid sensors that recognize foreign and incorrectly localized or processed host DNA and RNA. Double-stranded (ds)DNA sensors include (cyclic GMP-AMP synthase) cGAS, DNA-dependent activator of IRFs (DAI), absent in melanoma 2 (AIM2), and DNA-dependent protein kinase (DNA-PK), and activation of these receptors initiates various signaling cascades to generate an anti-microbial gene expression response ^2–6^. cGAS is the primary DNA sensor in most cell types and restricts a variety of pathogenic infections including viruses, bacteria, and fungi ^7,8^. Following DNA binding in a sequence-independent manner, cGAS synthesizes the second messenger 2’3’ cyclic GMP-AMP (cGAMP) which binds the protein stimulator of interferon genes (STING) on the endoplasmic reticulum ^9–13^. STING translocates to the Golgi and recruits TANK-binding kinase 1 (TBK1), which in turn activates the transcription factor complexes interferon regulatory factor (IRF) and nuclear factor kappa-light-chain-enhancer of activated B cells (NF-kB), inducing IRF and NF-kB nuclear translocation to stimulate IFN and pro-inflammatory cytokine expression, respectively.

Although the cGAS-STING pathway is canonically investigated in the context of infection, recent findings demonstrate it can be activated by host DNA to impact cellular homeostasis independent of pathogens. Tight regulation of cGAS and endogenous dsDNA localization is essential for preventing cGAS activation of an aberrant immune response that contributes to various diseases including autoimmune diseases such as systemic lupus erythematosus (SLE) and Aicardi-Goutieres syndrome (AGS) ^14–18^. Several cGAS regulatory mechanisms have been identified that inhibit the cGAS-STING pathway including protein-protein interaction with Beclin-1 to limit 2’3’ cGAMP production and deSUMOylation by SENP2 to induce cGAS degradation ^19,20^. Furthermore, cGAS serine phosphorylation is a major inhibitory mechanism that restricts cGAS recognition of host DNA. Akt phosphorylation of cGAS reduces its enzymatic function, while phosphorylation by CDK1-cyclin B restricts cGAS activity during mitotic entry, and Aurora kinase B hyperphosphorylation of cGAS prevents recognition of chromatin during mitosis ^21–23^. As cGAS is associated with numerous diseases, the identification of other cGAS regulatory mechanisms will provide the foundation for developing novel therapeutics.

cGAS stimulation is further associated with the induction of anti-tumor immunity in a variety of cancer subtypes by inducing expression of inflammatory genes and secretion of their protein products to recruit immune cells while also inducing replicative senescence that collectively restrict tumor growth and promote tumor clearance ^24,25^. Indeed, tumor-derived 2’3’ cGAMP is transferred to myeloid and B cells to induce NK cell-mediated tumor cell killing ^26^. Furthermore pharmacological activation of the cGAS-STING pathway induces an anti-tumor response suggesting that targeting cGAS in chemotherapeutic regimens would enhance patient outcomes ^27–30^. Unfortunately cGAS and STING are inhibited or downregulated in various cancers through epigenetic, hypoxia, and post-translational modification mechanisms ^31–33^. As cGAS-STING activation generates an anti-tumor immune response, we hypothesized that cancer-associated proteins hinder cGAS-STING signaling by inhibiting cGAS activity to promote cancer progression. The proto-oncogene non-receptor tyrosine kinase c-Src is involved in several cellular pathways that enhance cancer development including differentiation, proliferation, survival, and motility ^34^. Importantly c-Src expression and kinase activity are frequently elevated in various cancers including lung, breast, and colon cancer, and correlates with increased tumor initiation, progression, and metastasis ^35^. As such c-Src is pharmacologically inhibited by FDA-approved ATP competitive inhibitors such as bosutinib and dasatinib in chemotherapeutic cocktails for hematological cancer ^36^. As cGAS is frequently inhibited in cancer, further investigation is needed to identify cancer-associated proteins such as c-Src that prevent cGAS activation and are amenable to therapeutic intervention to drive an anti-tumor immune response.

c-Src phosphorylates a variety of intracellular proteins and as c-Src is highly expressed in cancer, it is likely that c-Src also impacts other pathways that are involved in cancer biology such as nucleic acid sensing. Therefore, we sought to investigate whether c-Src restricts cGAS activity through phosphorylation to limit an immune response. Here, we use publicly available data from The Cancer Genome Atlas (TCGA) to demonstrate that high c-Src expression correlates with reduced immune gene expression in various tumors, and that high c-Src and low cGAS expression also correlates with worse lung cancer patient survival. Furthermore, c-Src restricts a dsDNA-dependent IFN and NF-|B immune response in lung adenocarcinoma, acute monocytic leukemia, and immortalized kidney cell lines. c-Src inhibition enhances a cGAS-dependent immune response while c-Src overexpression inhibits it in a c-Src-kinase dependent manner. We further demonstrate that cGAS and c-Src interact with each other in cells and *in vitro* while c-Src restricts cGAS enzymatic activity to synthesize 2’3’ cGAMP potentially due to reduced cGAS DNA binding. Finally, we show that c-Src directly phosphorylates cGAS and genetically preventing phosphorylation at cGAS Y248 partially reduces the ability of c-Src to restrict cGAS signaling. Collectively, our study describes how the proto-oncogene c-Src regulates the DNA sensor cGAS and demonstrates how cancer-associated proteins can impact anti-tumor innate immune sensing.

## Results

### c-Src expression correlates with reduced immune signatures and patient survival in lung cancer

c-Src is frequently highly expressed in certain cancers and is associated with enhanced cancer development ^35^. In addition, anti-tumor cGAS-STING signaling is typically reduced or inhibited in cancer to promote tumor progression ^31–33^. Therefore, we hypothesized that cancer types with elevated c-Src expression levels would correlate with reduced innate immune gene expression due to inhibited cGAS DNA sensing. To first test this hypothesis, we used the Human Protein Atlas database to identify cancer types with high to medium c-Src expression based on immunohistochemistry (proteinatlas.org) ^37^. ∼90% of melanoma, ∼60% of bladder cancer, and ∼50% of lung adenocarcinoma patients are in this category while ∼0% of lymphoma patients express high to medium c-Src, demonstrating that not all cancers have high c-Src levels (Figure 1A). Next, we analyzed gene expression in melanoma, bladder cancer, and lung adenocarcinoma tumors using publicly available patient data in the human Cancer Genome Atlas (TCGA) with the UCSC Xena program ^38^. Interestingly, gene expression analysis revealed that high c-Src levels correlate with reduced expression of various immune genes, including IFN ®, the ISGs IFIT2 and PKR, and the NF-|B-transcribed gene IL6 in all three cancer types (Figure 1B-E; Figure S1A). This negative correlation is not true for all genes as expression of the RNA binding protein FUS varies in the different cancers. Importantly, other oncogenes do not appear to restrict an immune response as B-Raf and K-RAS expression, oncogenes that have increased activity in melanoma and lung adenocarcinoma, respectively, do not correlate with reduced immune gene expression (Figure S1B, C) ^39,40^. This data demonstrates that c-Src may regulate cGAS sensing in various cancers as a means of inhibiting an anti-tumor immune response.

**Figure 1:**
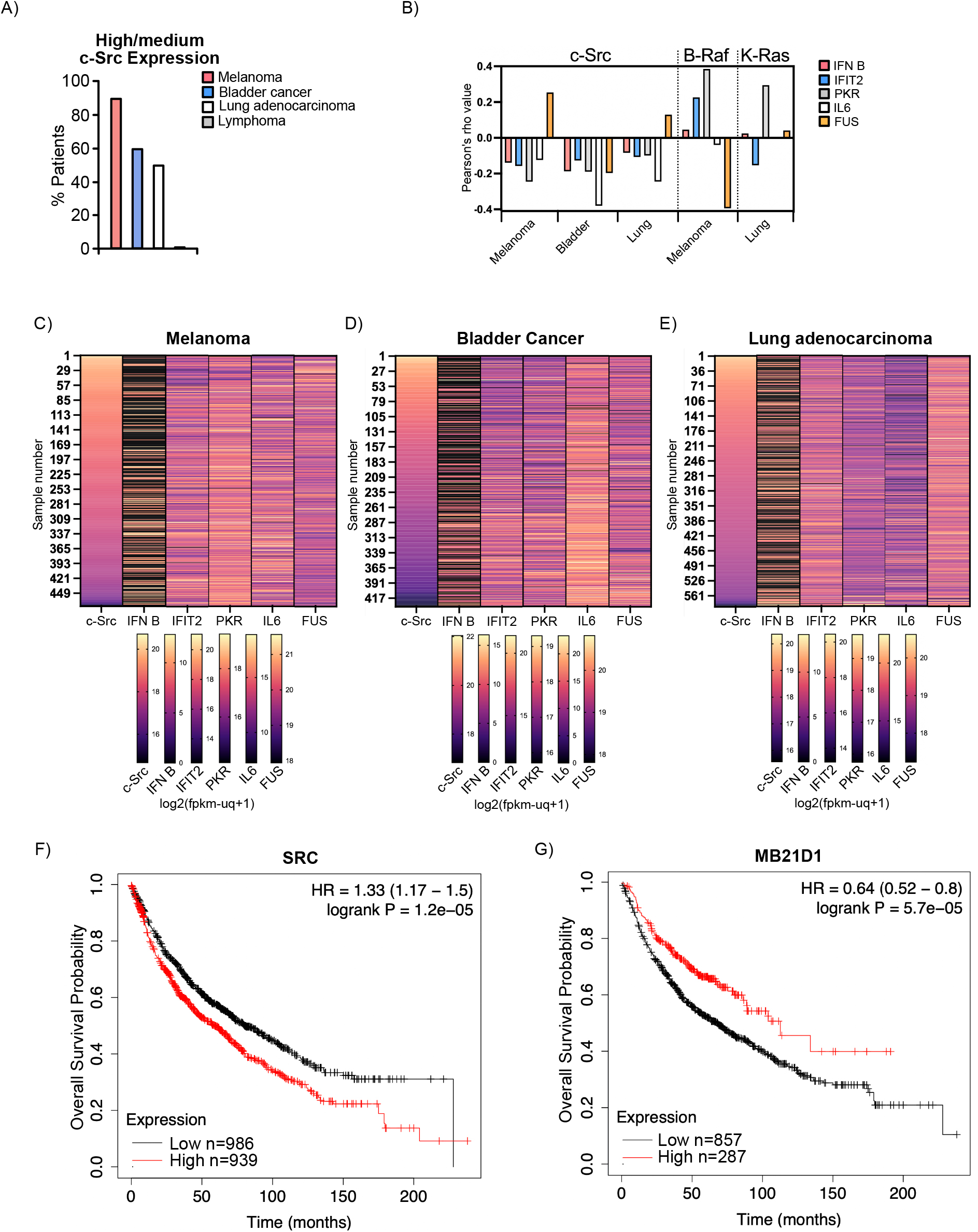
Reduced tumor innate immune gene expression and patient survival correlates with elevated c-Src expression in various cancer types. **(A)** Percent of patients with high/medium c-Src expression in different types of cancer based on immunohistochemistry. Data was analyzed through Human Protein Atlas. **(B)** Pearson’s rho correlation values for heat maps. **(C)** Heat map of TCGA data of c-Src, IFN ®, IFIT2, PKR, IL6, and FUS expression levels in melanoma patients. Data was analyzed through the UCSC Xena program. p values are in Figure S1. Expression range and corresponding colors are included for each gene representing log2(fpkm-uq+1). **(D)** Heat map of TCGA data for bladder cancer patients similar to (B). **(E)** Heat map of TCGA data for lung adenocarcinoma patients similar to (B). **(F)** Kaplan-Meier survival curves of lung cancer patients and c-Src (SRC) expression levels. Curves were plotted using KM Plotter. Expression cutoff was dictated by auto select best cutoff in program. FDR= 1%. HR, hazard ratio. Statistical significance was determined using the log-rank test. **(G)** Kaplan-Meier survival curves of lung cancer patients and cGAS (MB21D1) expression levels. Curves were plotted using KM Plotter. FDR= 3%. Also see Figure S1.

Given that elevated c-Src expression would inhibit cGAS signaling and reduce an anti-tumor immune response, we hypothesized that high c-Src levels correlate with reduced patient survival. To test this hypothesis, we plotted Kaplan–Meier survival curves using publicly available lung cancer patient data and analyzed c-Src expression levels. Kaplan–Meier analysis revealed that high c-Src levels strongly correlate with reduced survival (Figure 1F). We further determined that low cGAS expression also correlated with enhanced death, suggesting that loss of cGAS activity significantly diminishes patient outcomes (Figure 1G).

### c-Src restricts a cGAS-dependent dsDNA immune response in various cell types

c-Src phosphorylates a variety of proteins to impact different pathways in the cell. To determine if c-Src regulates a cGAS-dependent DNA sensing immune response, we first validated a cellular co-culturing method to analyze cGAS signaling without potential interference from the effects of c-Src overexpression in cells. Here the cyclic dinucleotide 2’3’ cGAMP produced in donor cells is transferred to recipient L929-IFN-stimulated response element (ISRE)--luciferase (LUC) reporter cells via connexins in gap junctions to induce ISRE-LUC expression ^41,42^. We verified that various cyclic dinucleotides including 2’3’ cGAMP are transferred from HEK293T cells transfected with different cyclic dinucleotide cyclases into L929-ISRE-LUC cells by the induction of an ISRE response (Figure S2A). Furthermore, transfection of cyclic dinucleotides directly into L929-ISRE-LUC cells induced ISRE activity, demonstrating that they are immunostimulatory (Figure S2B). We also validated that 2’3’ cGAMP is required to induce an ISRE response by transfecting HEK293T cells with cGAS and Poxin, a known cGAMP cleavage enzyme, and observed a loss of ISRE induction (Figure S2C) ^43^. Finally, we confirmed that 2’3’ cGAMP is transferred via gap junctions by treating the co-cultured cells with meclofenamic acid (MFA), a gap junction antagonist, and did not detect an ISRE response (Figure S2D). Overall, these data demonstrate that 2’3’ cGAMP is transferred from DNA transfected cells into co-cultured L929-ISRE-LUC cells to induce ISRE activity and can be used to determine if c-Src regulates a cGAS-dependent immune response.

A549 lung adenocarcinoma cells express cGAS and high levels of c-Src compared to HEK293T and monocytic THP-1 cells, making them an ideal cell line to investigate c-Src regulation of cGAS (Figure S3A). To determine if c-Src regulates cGAS signaling, we treated A549 cells with the c-Src small molecule inhibitor dasatinib. Importantly dasatinib treatment inhibited c-Src activity in A549 cells as autophosphorylation at c-Src Y416 is lost after treatment (Figure S3B) ^44^. Following transfection of the cGAS ligands herring testes DNA (HT-DNA) and interferon stimulatory DNA (ISD45), and co-culturing with L929-ISRE-LUC cells, A549 cells treated with dasatinib induced significantly higher ISRE activity as compared to mock treated cells, suggesting that c-Src inhibits cGAS signaling (Figure 2A). We further verified the increase in ISRE activity is not due to dasatinib off-target effects as saracatinib treatment, a different c-Src inhibitor, also increased DNA sensing in A549 cells (Figure S3C).

**Figure 2:**
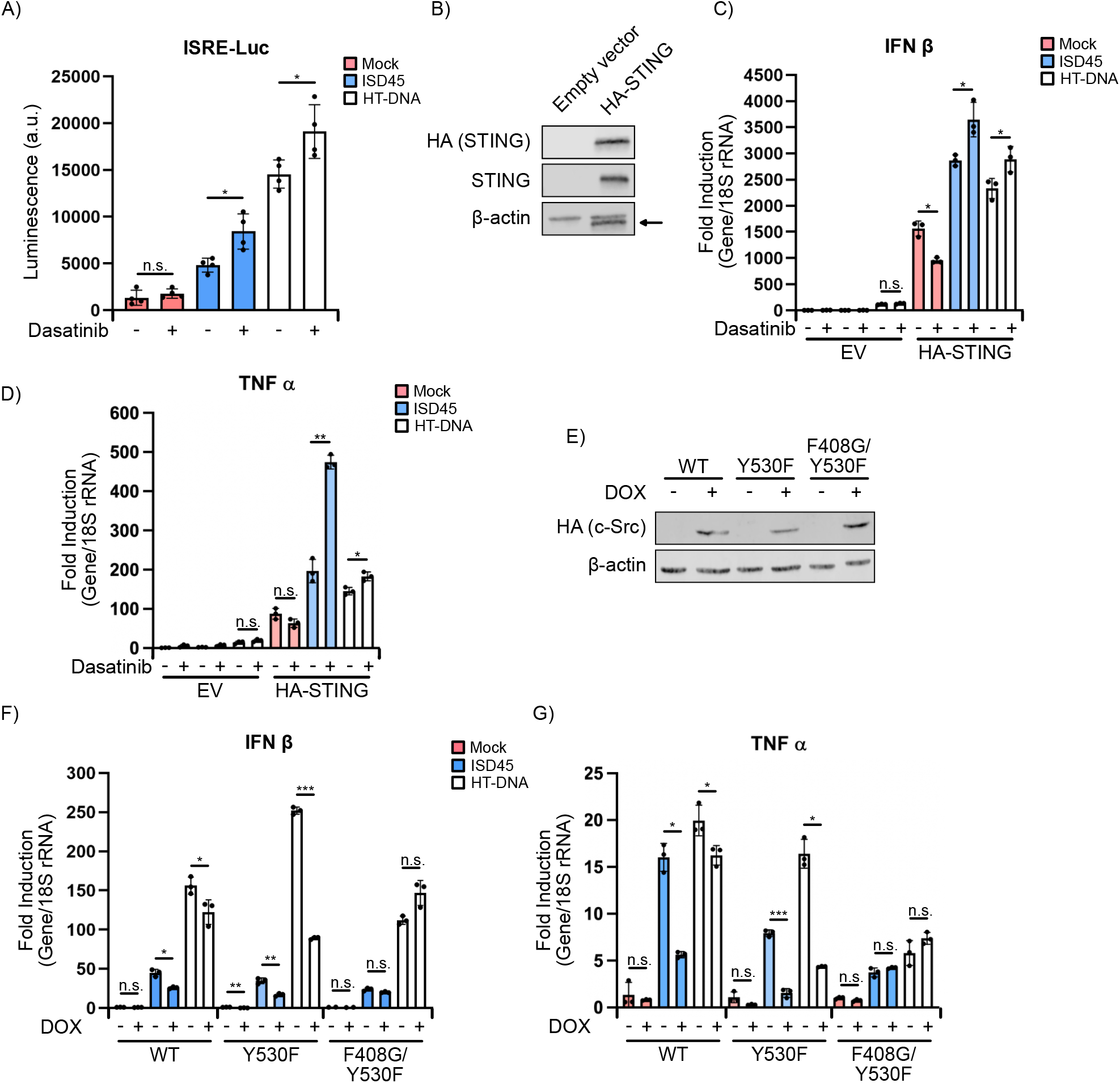
c-Src restricts a DNA sensing immune response. **(A)** Co-culture of L929-ISRE-LUC cells with A549 cells treated with vehicle or dasatinib (30nM) for 48 h followed by transfection with indicated DNA ligands for 6 h. **(B)** Western blot of A549 cells transfected with pcDNA3 empty vector (EV) or HA-STING pcDNA3. Arrow depicts not-specific band. ®-actin was used as a loading control. **(C)** RT-qPCR analysis of IFN ® levels from cells in (B) treated with vehicle or dasatinib for 48 h followed by transfection with indicated DNA ligands for 6 h. All samples were normalized to 18S rRNA. **(D)** RT-qPCR of TNF ⟨ levels from cells in (C). **(E)** Western blot of THP-1 WT and mutant HA-c-Src-tet-RB SB cells treated with 1 μg/ml doxycycline (DOX) for 48 h. ®-actin was used as a loading control. **(F)** RT-qPCR analysis of IFN ® levels from cells in (E) treated with vehicle or DOX for 48 h followed by transfection with indicated DNA ligands for 6 h. All samples were normalized to 18S rRNA. **(G)** RT-qPCR of TNF ⟨ levels from cells in (F). *n*= 4 biological replicates for luciferase assay, and *n*= 3 biological replicates and representative qPCR with technical triplicates is shown. Also see Figure S2, S3.

Given that A549 cells do not express STING we next sought to determine whether STING expression would permit functional cGAS signaling and increased immune gene induction following c-Src inhibition independent of a co-culture. A549 cells were treated with dasatinib and transfected with a HA-tagged STING expression vector prior to cGAS ligand transfection and RNA isolation (Figure 2B). Importantly STING expression increased IFN β, a type I interferon, mRNA levels in mock treated cells as the vector itself activated cGAS, confirming the cGAS-STING signaling pathway is complete (Figure 2C). c-Src inhibition induced significantly higher expression of various immune genes including IFN β, the pro-inflammatory gene TNF ⟨, and the interferon stimulated gene (ISG) IFIT2 following ligand transfection, demonstrating c-Src inhibition enhances a STING-dependent IFN and NF-|B immune response (Figure 2D; Figure S3D).

Next, we sought to determine whether c-Src overexpression impacts DNA signaling. As transient transfection of DNA vectors induce cGAS activity, we generated stable doxycycline (DOX) inducible HA-c-Src Sleeping Beauty (SB) vectors that insert into the genome following selection ^45^. Wild-type (WT), kinase dead (KD) (K298M and F408G/Y530F), and kinase constituently active (Y530F) c-Src mutants were produced to determine if c-Src kinase activity is responsible for inhibiting DNA sensing ^46–50^. Fascinatingly, although the WT and c-Src mutants were induced by DOX treatment in A549 cells, ISRE activity was not impacted following cGAS ligand transfection (Figure S3E, F). It is possible the lack of ISRE induction is due to high levels of endogenous c-Src in A549 cells where cGAS inhibition is already saturated. Therefore, we used THP-1 cells to overexpress HA-c-Src as they contain a functional cGAS-STING pathway and minimal c-Src (Figure 2E). Importantly overexpression of WT and Y530F c-Src in THP-1 cells significantly reduced IFN β and TNF ⟨ induction following cGAS ligand transfection while the F408G/Y530F kinase dead mutant did not impact their expression, suggesting c-Src overexpression restricts DNA sensing in a kinase-dependent manner (Figure 2F, G). Furthermore, dasatinib treatment of THP-1 cells followed by cGAS ligand transfection did not impact IFN β expression, further demonstrating that c-Src inhibition of cGAS depends on c-Src and cGAS expression levels as c-Src is minimally expressed in THP-1 cells (Figure S3G).

HEK293T cells do not express cGAS but express detectable c-Src, allowing us to transfect DNA expression plasmids without activating an immune response. Therefore, to provide additional evidence that c-Src overexpression inhibits cGAS signaling, we transfected HEK293T cells with WT, KD, and constituently active HA-c-Src mutants with or without cGAS followed by co-culturing with L929-ISRE-LUC cells (Figure 3A). Consistent with c-Src overexpression restricting cGAS signaling in THP-1 cells, WT and Y530F c-Src significantly reduced ISRE activity in a cGAS-dependent manner (Figure 3B). Importantly both the K298M and F408G/Y530F KD mutants did not impact DNA sensing, validating a kinase-dependent inhibition. We further analyzed murine IFN β mRNA levels in the co-cultured L929-ISRE-LUC cells as a different immune response measurement. Similar to ISRE-LUC induction, c-Src restricted IFN β expression in a kinase-dependent manner (Figure 3C). c-Src kinase activity further restricted cGAS-dependent NF-|B signaling as demonstrated through a NF-|B reporter luciferase assay (Figure S4A).

**Figure 3:**
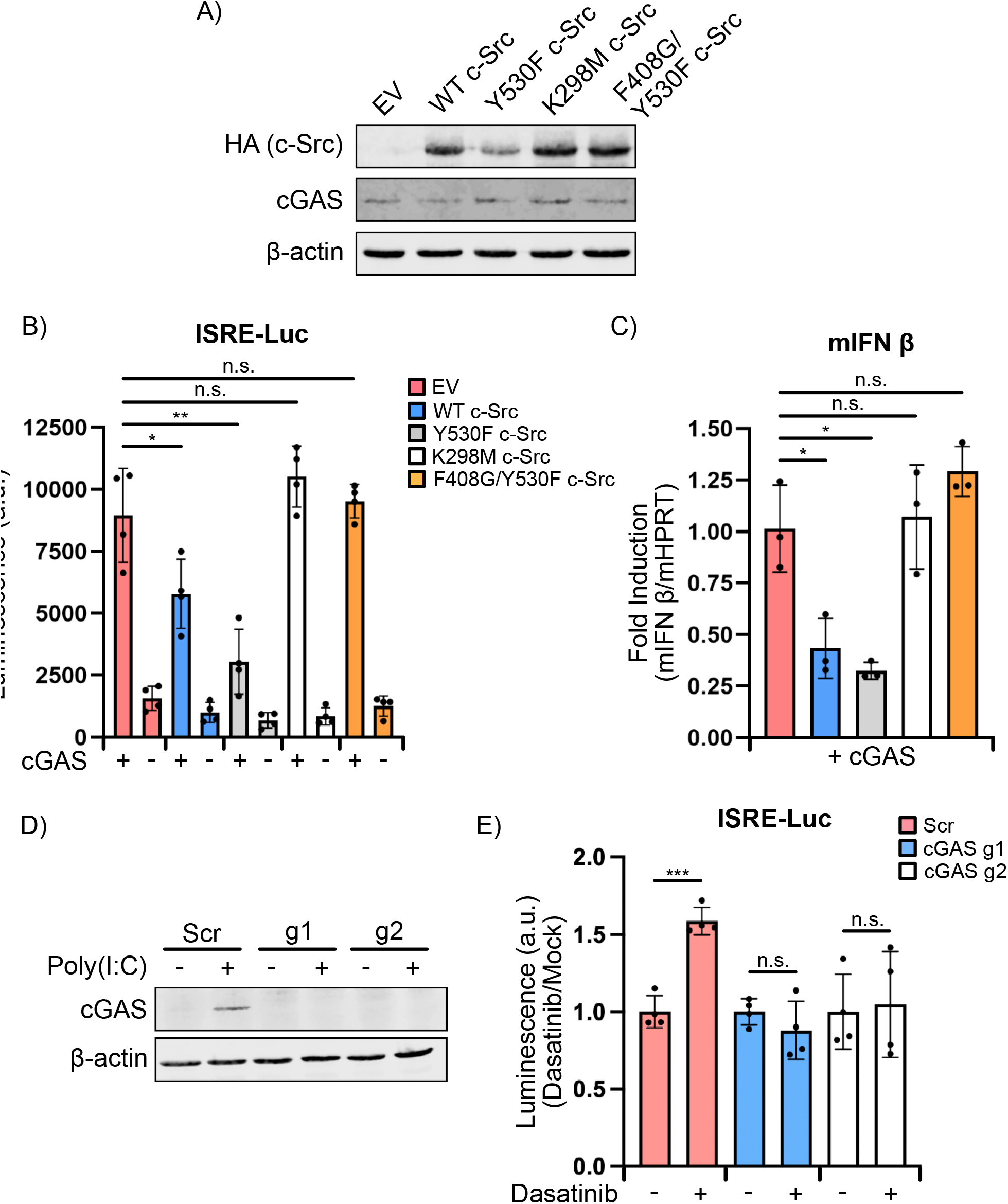
cGAS-dependent immune responses are hindered by c-Src. **(A)** Western blot of HEK293T cells transfected with 75 ng cGAS-pcDNA3 and 925 ng of EV, WT c-Src-HA pcDNA3, or mutant c-Src-HA pcDNA3 for 24 h. ®-actin was used as a loading control. **(B)** Co-culture of L929-ISRE-LUC cells with cells in (A). **(C)** RT-qPCR analysis of murine IFN ® levels from co-cultured L929-ISRE-LUC cells in (B). All samples were normalized to mHPRT. **(D)** Western blot of A549 cGAS KO cells. Non-target (Scr) guide RNA and two independent cGAS guide RNAs are shown. cGAS expression was induced by transfecting 1 μg/ml poly(I:C) for 16 h. ®-actin was used as a loading control. **(E)** Co-culture of L929-ISRE-LUC cells with A549 Scr and cGAS KO cells treated with vehicle (mock) or dasatinib (30nM) for 48 h followed by transfection with HT-DNA for 6 h. Mock luciferase levels for each cell line was set to 1. *n*= 4 biological replicates for luciferase assay, and *n*= 3 biological replicates and representative qPCR with technical triplicates is shown. Also see Figure S4.

c-Src is a member of the Src family kinases (SFKs), a large family of conserved kinases that have redundant function and are inhibited by c-Src inhibitors, albeit with higher IC50s ^51^. Therefore, we sought to determine whether c-Src depletion also enhances cGAS activity by generating DOX-inducible short hairpin (sh)RNAs that target c-Src in HEK293T cells. c-Src mRNA levels were reduced by ∼90% of a non-target shRNA using two independent c-Src shRNAs (Figure S4B). c-Src depletion significantly amplified cGAS-dependent ISRE activity following ISD45 transfection and co-culturing with L929-ISRE-LUC cells, demonstrating that c-Src inhibition and depletion increases cGAS signaling (Figure S4C). Importantly, c-Src expression did not impact ISRE activity in response to 2’3’ cGAMP transfection, suggesting that inhibition occurs upstream of STING in the pathway.

Lastly, to determine whether enhanced ISRE activity observed following c-Src inhibition and co-culturing is cGAS-dependent, we generated cGAS CRISPR/Cas9 knockouts (KO) in A549 cells. Given that cGAS is expressed at low levels under basil conditions and is an ISG, A549 cGAS KO cells were transfected with the synthetic double-stranded RNA analog poly(I:C) to induce expression through the RIG-I-like receptors (RLR) RNA signaling pathway followed by Western blotting ^52^. cGAS was completely knocked out using two independent guide RNAs as compared to a non-target scramble (Scr) guide and we further validated the KOs by identifying the CRISPR/Cas9 induced mutations (Figure 3D). To determine whether c-Src inhibits DNA sensing in a cGAS-dependent manner, we treated WT and KO cells with dasatinib followed by cGAS ligand transfection and co-culturing. While dasatinib treatment enhanced ISRE activity in WT cells, the increase was completely abolished in the cGAS KO cells (Figure 3E). Collectively, these data demonstrate that c-Src regulates a cGAS-dependent interferon and NF-|B gene expression response in multiple cell types.

### c-Src interacts with cGAS and inhibits its enzymatic activity

Next, we investigated whether cGAS and c-Src interact with each other which could suggest a direct inhibitory mechanism. To determine if the proteins interact in cells, HEK293T cells were transfected with equal concentrations of empty vector (EV), FLAG-cGAS, or c-Src-HA expression plasmids for 24 h, lysed and subjected to a FLAG-immunoprecipitation (IP), and proteins were analyzed by Western blotting. cGAS was efficiently IP’d with and without c-Src co-expression while c-Src, but not the loading control ®-actin, was pulled-down in the IP (Figure 4A). Moreover, the reciprocal HA-IP revealed FLAG-cGAS was also co-IP’d with c-Src-HA (Figure 4B). To determine whether cGAS and c-Src interact in THP-1 cells, which is a more physiologically relevant cell type, we treated inducible HA-c-Src SB THP-1 cells with DOX to induce HA-c-Src expression followed by a HA-IP. Importantly endogenous cGAS was co-IP’d with c-Src, demonstrating that cGAS and c-Src interact in various cell types (Figure 4C).

**Figure 4:**
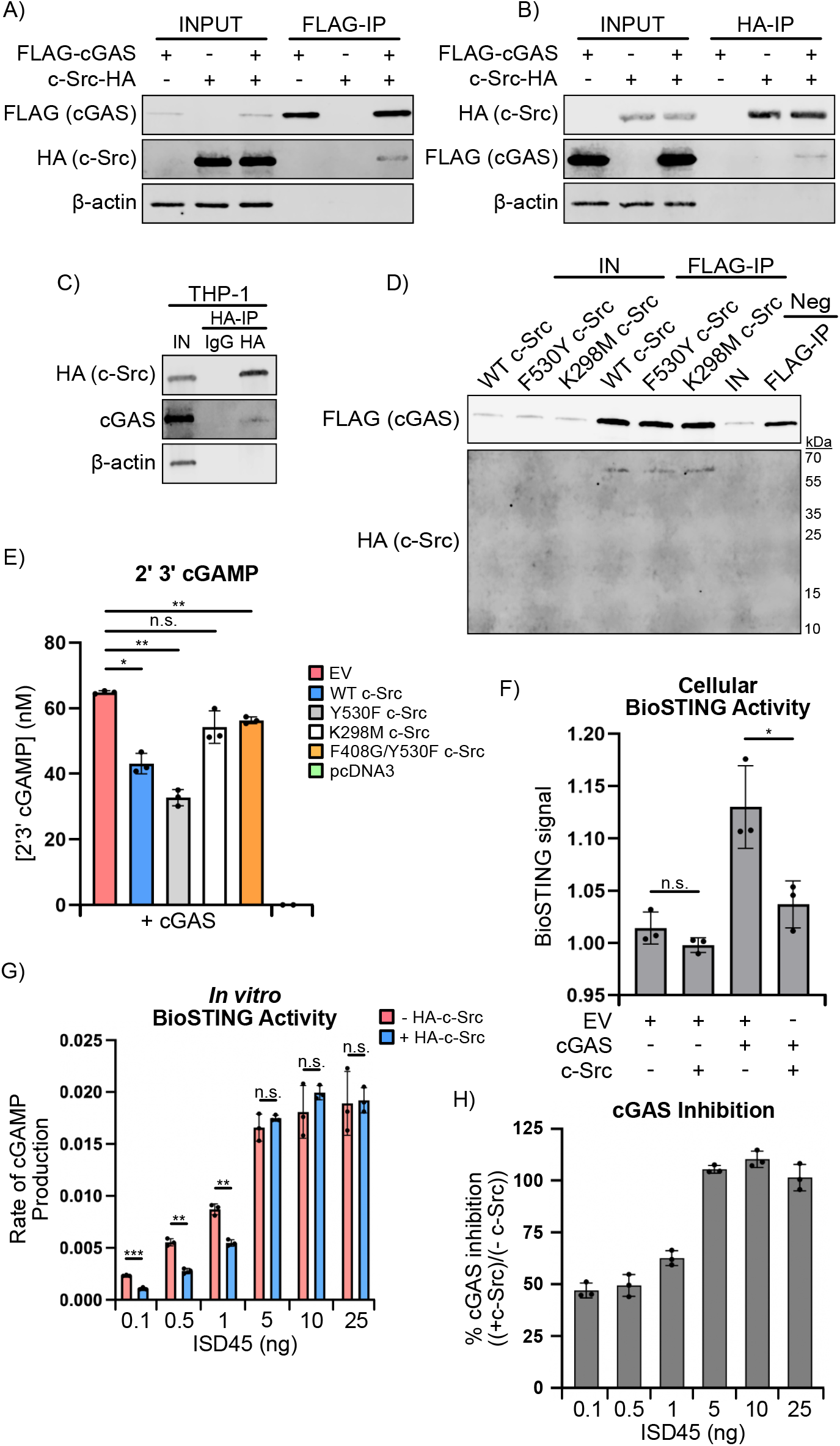
c-Src directly interacts with cGAS and inhibits cGAS activation. **(A)** Western blot of HEK293T cells transfected with equal concentrations of EV, FLAG-cGAS pcDNA3, or c-Src-HA pcDNA3 for 24h followed by FLAG immunoprecipitation (IP). ®-actin was used as a loading control. **(B)** Similar to (A) but HA IP. **(C)** Western blot of THP-1 WT HA-c-Src pSB-tet-RB SB cells treated with DOX for 48 h and poly(I:C) for 16 h followed by a HA IP. ®-actin was used as a loading control. **(D)** Western blot of *in vitro* co-IP of FLAG-cGAS, and WT or mutant HA-c-Src. Equal concentrations of cGAS and c-Src were incubated together followed by FLAG IP. A control HA-tagged protein (12 kDa) was used as a negative control. **(E)** 2’3’ cGAMP EIA of HEK293T cells transfected with cGAS-pcDNA3, and EV, WT, or mutant c-Src-HA pcDNA3 for 24 h. pcDNA3 alone was used as a negative control. **(F)** BioSTING FRET assay from HEK293T cells transfected with EV, cGAS pcDNA3, and c-Src-HA pcDNA3 for 24 h prior to cell lysis. Lysates were incubated with purified BioSTING and BioSTING activity was analyzed by FRET signal at 5 min post incubation. **(G)** BioSTING FRET assay from purified FLAG-cGAS that had been phosphorylated by WT HA-c-Src or mock treated prior to incubation with BioSTING and various concentrations of ISD45. BioSTING activity was plotted by analyzing rate of 2’3’ cGAMP production as it relates to ISD45 concentration. **(H)** Percentage of cGAS inhibition by c-Src from data in (G). Mock treatment was set at 1 and HA-c-Src treatment was compared to mock. *n*= 3 biological replicates and representative assays with technical triplicates are shown. Also see Figure S5.

A direct interaction is required for c-Src to phosphorylate cGAS. Therefore, to investigate whether cGAS and c-Src directly interact, we purified FLAG-cGAS and HA-c-Src from *E. coli* and HEK293T cells, respectively. We first validated that the purified proteins were active. FLAG-cGAS was incubated with ISD45, ATP, GTP, and purified BioSTING, a STING-based Förster resonance energy transfer (FRET) biosensor which measures 2’3’ cGAMP abundance by FRET signal ^53^ (Figure S5A). Increasing concentrations of FLAG-cGAS enhanced BioSTING activity temporally due to 2’3’ cGAMP production, demonstrating that FLAG-cGAS is functional (Figure S5B). Purified WT, K298M, and Y530Y HA-c-Src kinase activity were analyzed by *in vitro* kinase assays followed by Western blotting for autophosphorylation at Y416 (Figure S5C). Importantly WT and Y530F were phosphorylated but K298M was not, confirming HA-c-Src is active (Figure S5D). To determine if cGAS directly interacts with HA-c-Src and if the interaction is impacted by c-Src kinase activity, equal concentrations of FLAG-cGAS and HA-c-Src were incubated together followed by a FLAG IP and analysis by Western blotting. Fascinatingly WT, Y530F, and K298M c-Src all interact with cGAS while a HA-tagged negative control protein (12 kDa) did not interact (Figure 4D). Overall, these results suggest that c-Src directly interacts with cGAS, and kinase activity does not affect binding.

As c-Src restricts cGAS signaling in a kinase-dependent manner we hypothesized that c-Src limits cGAS enzymatic ability to synthesize 2’3’ cGAMP, resulting in a reduced immune response. To determine if c-Src restricts 2’3’ cGAMP production, we perform a 2’3’ cGAMP enzyme immunoassay (EIA) on HEK293T cells expressing cGAS and the various c-Src-HA constructs. Interestingly 2’3’ cGAMP levels were significantly reduced by WT and Y530F c-Src but not the KD mutants (Figure 4E). We further analyzed 2’3’ cGAMP production by incubating cell lysates from HEK293T cells expressing cGAS and WT c-Src-HA with purified BioSTING followed by FRET analysis. BioSTING was activated in a cGAS-dependent manner and activity was significantly reduced with c-Src co-expression (Figure 4F). Finally, we determined whether c-Src restricts cGAS activity *in vitro* by first subjecting purified FLAG-cGAS to an *in vitro* kinase assay using HA-c-Src, followed by determination of 2’3 cGAMP production with BioSTING at various concentrations of ISD45. c-Src significantly reduced the rate of 2’3’ cGAMP production in a DNA concentration-dependent manner (Figure 4G). At low DNA concentrations, c-Src exhibited ∼50% inhibition, while 2’3’ cGAMP production was indistinguishable at elevated DNA concentrations (Figure 4H). Analysis of the DNA required to achieve 50% of maximum cGAS activity revealed a 2-fold shift from 1.2 to 2.4 ng of ISD45. These data demonstrate that c-Src interacts with cGAS, and c-Src restricts cGAS enzymatic activity in a kinase-dependent manner, potentially through an altered affinity of cGAS for activating DNA.

### cGAS-DNA binding is reduced by c-Src kinase activity

Inhibition of cGAS activity can occur through a variety of mechanisms. Restricting cGAS DNA binding is one such process and has frequently been identified as a means to limit cGAS signaling. cGAS phosphorylation by Aurora kinase B prevents chromatin binding during mitosis while BAF restricts nuclear cGAS activity by outcompeting cGAS for DNA binding ^23,54^. To validate the *in vitro* kinetic observations and investigate if c-Src reduces cGAS DNA binding, we transfected HEK293T cells with FLAG-cGAS with or without c-Src-HA, crosslinked cells, and performed a FLAG-IP coupled with qPCR (Figure 5A). As plasmid DNA is a cGAS ligand we analyzed bound pcDNA3, the plasmid backbone for the expression vectors, by qPCR at three different locations on the plasmid. Importantly the FLAG IP enriched cGAS at similar efficiency with and without c-Src co-expression by Western blot (Figure 5B). IP-qPCR percent input analysis revealed that FLAG-cGAS expression increased DNA binding as compared to mock treated cells (Figure 5C). Intriguingly, c-Src co-expression significantly reduced cGAS DNA binding at three locations on the plasmid as compared to cGAS expression alone. cGAS binding is unique to the transfected plasmids as host GAPDH is not bound to cGAS under any condition. We further determined that cGAS DNA binding is reduced in a c-Src kinase-dependent manner by performing a FLAG-cGAS IP-qPCR with the K298M KD mutant and observed a rescue in DNA binding similar to EV levels (Figure 5D).

**Figure 5:**
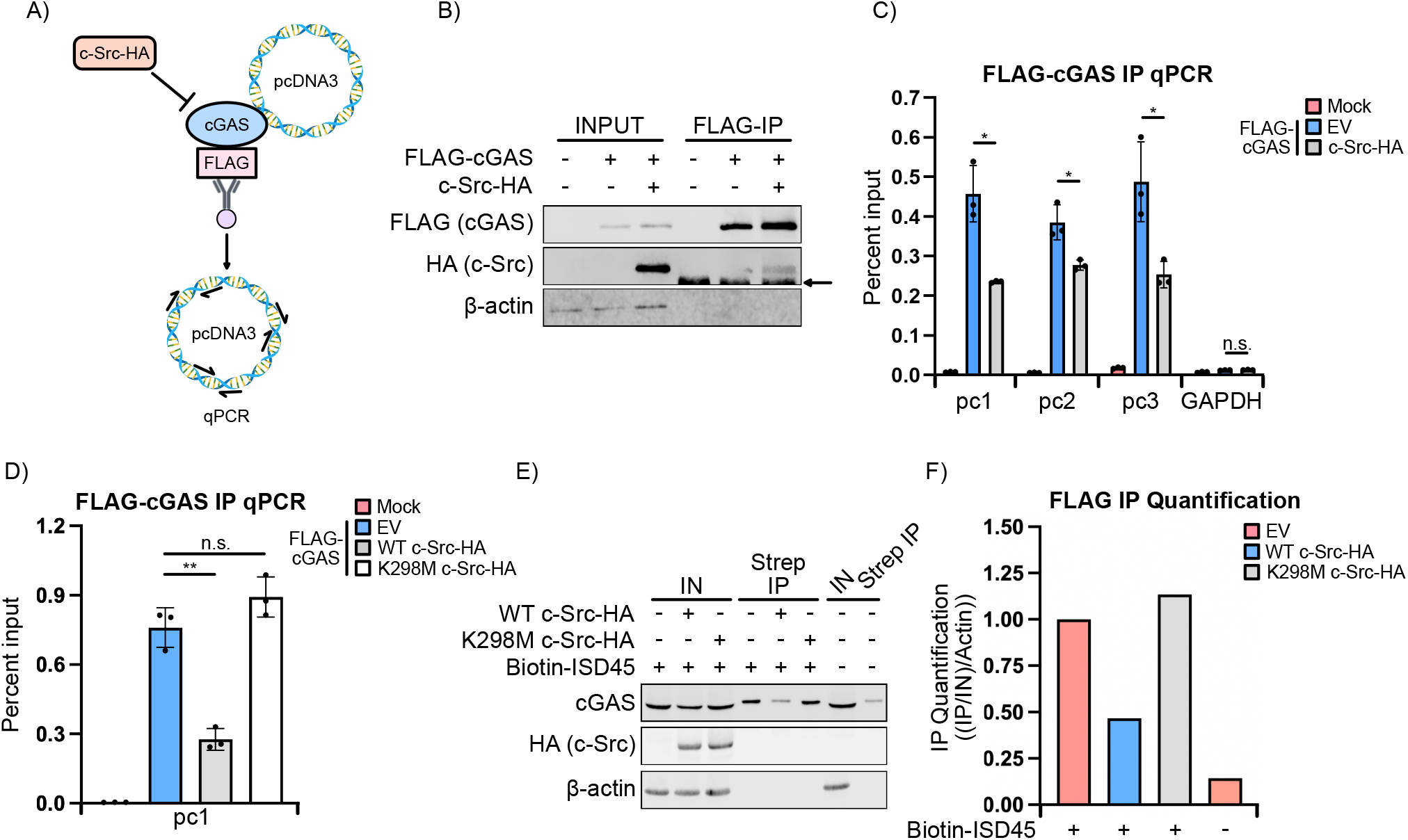
c-Src kinase activity restricts cGAS DNA binding. **(A)** Schematic of FLAG-cGAS IP-qPCR. HEK293T cells were transfected with EV, FLAG-cGAS pcDNA3, and c-Src-HA pcDNA3 for 24 h. Cells were crosslinked and subjected to FLAG IP coupled with qPCR analysis to detect bound pcDNA3 vector. **(B)** Western blot of FLAG-cGAS IP-qPCR described in (A). Arrow depicts not-specific band. ®-actin was used as a loading control. **(C)** IP-qPCR analysis from FLAG-IP in (B). Samples were quantified by percent input. Three different regions of pcDNA3 (pc1, pc2, pc3) were analyzed as well as non-specific GAPDH. **(D)** IP-qPCR analysis of HEK293T cells transfected with FLAG-cGAS pcDNA3, and EV, WT, or K298M c-Src-HA pcDNA3 followed by FLAG-cGAS-IP. **(E)** Western blot of HEK293T cells transfected with FLAG-cGAS pcDNA3, and EV, WT, or K298M c-Src-HA pcDNA3 for 24 h. Cells were then transfected with 4 μg/ml biotinylated or unmodified ISD45 for 2 h followed by streptavidin (strep) IP. ®-actin was used as a loading control. **(F)** Quantification of FLAG-cGAS enrichment from (E). *n*= 3 biological replicates and representative assays with technical triplicates are shown.

Lastly, to further examine c-Src restriction of cGAS DNA binding we isolated transfected cGAS ligands during c-Src overexpression and analyzed bound cGAS. HEK293T cells expressing FLAG-cGAS with or without c-Src-HA were transfected with biotinylated ISD45 and subjected to a streptavidin (strep) IP followed by Western blot analysis. Importantly the strep IP of biotinylated ISD45 enriched for cGAS over the non-modified ISD45 while c-Src reduced cGAS binding in a kinase-dependent manner (Figure 5E, F). Overall, these results demonstrate that c-Src kinase activity limits cGAS DNA binding to restrict cGAS function.

### cGAS is phosphorylated by c-Src

Given that c-Src directly interacts with cGAS, we sought to determine whether c-Src phosphorylates cGAS. *In vitro* kinase assays were performed on purified FLAG-cGAS by HA-c-Src and phosphorylation was analyzed by phosphor imaging. FLAG-cGAS and HA-c-Src are expressed at similar molecular weights so distinguishing between their phosphorylation status is difficult. Therefore, we varied the concentration of cGAS and c-Src added to the reaction. Comparable concentrations of c-Src incubated with increasing amounts of cGAS led to enhanced phosphorylation signal presumably on cGAS (Figure 6A). Furthermore, the phosphorylation signal intensified as increasing amounts of c-Src were used in a cGAS-dependent manner (Figure 6B). Finally, WT and Y530F c-Src phosphorylated cGAS while K298M did not, demonstrating that c-Src phosphorylates cGAS *in vitro* (Figure 6C).

**Figure 6:**
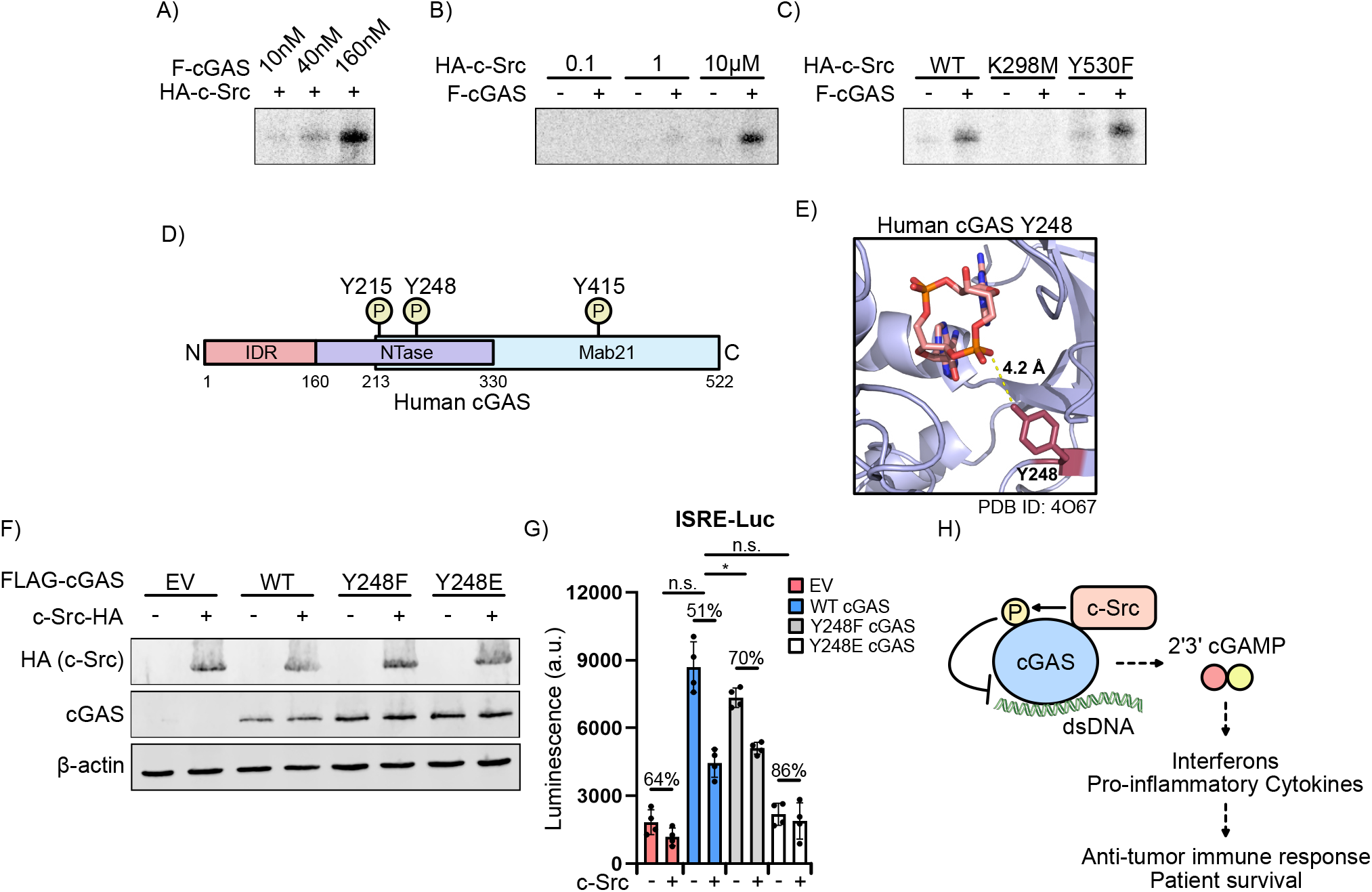
cGAS is phosphorylated by c-Src and cGAS Y248 is partially responsible for c-Src mediated inhibition. **(A)** *In vitro* kinase assay of 10 μM HA-c-Src with various concentrations of FLAG-cGAS. Phosphorylation was analyzed by phosphor imaging. **(B)** *In vitro* kinase assay similar to (A) of 160 nM FLAG-cGAS with various concentrations of HA-c-Src. **(C)** *In vitro* kinase assay similar to (A) of 160 nM FLAG-cGAS with 10 μM of WT and mutant HA-c-Src. **(D)** Schematic of human cGAS domains and observed tyrosine (Y) phosphorylation in PhosphoSitePlus. **(E)** Human cGAS structure (PDB ID: 4O67) with 2’3’ cGAMP in active site. Y248 is highlighted in red. **(F)** Western blot of HEK293T cells transfected with EV, WT, or Y248E/F FLAG-cGAS pcDNA3 mutants with or without c-Src-HA pcDNA3 for 24 h. ®-actin was used as a loading control. **(G)** Co-culture of L929-ISRE-LUC cells with cells in (F). **(H)** Model of c-Src inhibition of cGAS to restrict a DNA-dependent immune response and reduce innate immune gene expression in tumors. *n*= 4 biological replicates for luciferase assay, and *n*= 3 biological replicates other assays are shown.

To determine which cGAS tyrosine residues are phosphorylated by c-Src and their impact on cGAS activity, we first analyzed known and observed modified sites using the PhosphoSitePlus database (Figure 6D) ^55^. cGAS Y215 is in the DNA binding domain and phosphorylation promotes cytoplasmic localization to reduce a DNA damage-induced immune response as well as limits DNA binding ^56,57^. cGAS Y248 and Y415 phosphorylation have only been detected using proteomic discovery mass spectrometry and not verified. Interestingly Y248 is in the cGAS enzymatic pocket and in close proximity to synthesized 2’3’ cGAMP (Figure 6E)^58^. As cGAS Y215, Y248, and Y415 are phosphorylated residues, and Y215 and Y248 are in unique cGAS domains, we sought to determine what impact phosphorylation at these sites would have on c-Src restriction of cGAS signaling. Tyrosine residues were mutated to phenylalanine to prevent phosphorylation or glutamic acid, which functions as a phosphomimetic. To determine what affect the cGAS tyrosine mutants have on c-Src inhibition, HEK293T cells were transfected with WT or tyrosine mutant FLAG-cGAS with or without c-Src-HA followed by co-culturing with L929-ISRE-LUC cells. Interestingly only Y248 mutants were expressed at similar levels with or without c-Src, potentially due to reduced Y215 and Y415 mutant stability (Figure 6F). Therefore, we only investigated Y248 as reduced cGAS expression with c-Src would not permit accurate analysis. Fascinatingly, while c-Src again reduced WT cGAS signaling by ∼49%, Y248F was only inhibited ∼30%, suggesting that blocking Y248 phosphorylation restricts c-Src inhibition (Figure 6G). Furthermore, Y248E was not active under any conditions. Collectively, these results demonstrate that cGAS is phosphorylated by c-Src and that eliminating phosphorylation of cGAS Y248 partially blocks c-Src inhibition of cGAS signaling.

In summary, c-Src phosphorylates cGAS which inhibits cGAS DNA binding, 2’3’ cGAMP production, and an innate immune response while high c-Src expression correlates with reduced immune gene expression and patient survival in various cancer types (Figure 6H).

## Discussion

The cGAS-cGAMP-STING signaling pathway has garnered significant interest due to its pivotal roles during viral infection, autoimmunity, as well as the induction of anti-tumor immunity. Indeed, a hallmark of highly metastatic cancers is the presence of chromosomal instability, which results in the leakage of cytosolic dsDNA species through the formation of micronuclei ^59^. As such, tumor cells must employ a variety of strategies to dampen intrinsic cGAS-STING signaling in order to evade protective anti-tumor immune surveillance. Our results reveal a novel mechanism of tumor immune evasion through direct phosphorylation of cGAS by the prototypical oncoprotein c-Src. Moreover, using cancer cell lines and co-culture models, we identify c-Src inhibition as a promising strategy of augmenting cancer-cell intrinsic, as well as paracrine, innate immune activation in the context of cancer immunotherapy. Finally by leveraging publicly available cancer genomics data, we find that enhanced tumor c-Src expression is inversely correlated with type I interferon gene expression and associated with worse patient outcomes. Collectively, these data open up novel therapeutic avenues which could be leveraged to improve the immunogenicity of advanced and metastatic malignancies through the restoration of cancer-cell intrinsic cGAS signaling.

c-Src–the first and most extensively studied oncogene–is the founding member of a nine-gene family of non-receptor tyrosine kinases also known as SRC family kinases (SFKs), which are involved in the regulation of a variety of signaling processes. In the context of tumorigenesis, c-Src has been implicated in mitogenic signaling, cell-cycle progression, survival, angiogenesis, invasion, metastasis as well as resistance to chemotherapeutics. Indeed, aberrant c-Src activity has been identified in a variety of human malignancies including melanoma, lung cancers including non-small cell lung cancer, urothelial malignancies, and breast cancers, among many others. There is an urgent need for a more detailed understanding of the cellular targets and processes modulated by oncogenic c-Src signaling.

In recent years, strategies of augmenting STING signaling have been the subject of intense investigation for eliciting anti-tumor immune responses, especially for immunologically ‘cold’ tumors through the activation of tumor-resident myeloid cells and vasculature ^26,28,60,61^.

The clinical importance of STING-mediated anti-tumor immunity is perhaps best exemplified by non-muscle invasive bladder cancer where intravesical immunotherapy with the c-di-AMP producing bacteria, BCG, is a mainstay of treatment ^62,63^. In pre-clinical models of urothelial cancer, BCG strains engineered to overproduce c-di-AMP have yielded increased trained immunity, improved tumor clearance, and prolonged survival ^64^. Additionally, STING agonists including the synthetic, non-hydrolyzable cyclic dinucleotide analog, ADU-S100, are currently under investigation for a variety of solid organ and hematologic malignancies either as monotherapy or in combination with immune checkpoint therapies. On the other hand, indiscriminate STING hyperactivation in non-cancer cells may yield undesired off-target effects including vasculitides, which may limit the patient tolerability of these agents. Furthermore, recent findings demonstrate that systemic delivery of cyclic dinucleotides, including 2’3’ cGAMP, repress anti-tumor immunity due to (IL)-35^+^ regulatory B cell expansion that restricts NK cell-mediated tumor killing ^65^. Our findings suggest that c-Src inhibition could be leveraged to induce endogenous production of 2’3’ cGAMP in the tumor microenvironment (TME), which would eliminate the need for systemic treatment while still eliciting anti-tumor immunity.

Identification of cancer-cell specific strategies of cGAS-STING inhibition could therefore yield new therapeutic strategies for overcoming tumor-immune evasion and resistance to T-cell mediated immunotherapies. Indeed, repression of cGAS and STING expression through promoter hypermethylation has been observed across a variety of tumor types. To that end, pharmacological inhibition of DNA methylation has been proposed as one such strategy of restoring tumor cell intrinsic cGAS-STING signaling and downstream cytotoxic T-cell infiltration and killing ^66^. Similarly, upregulation of the cGAMP phosphodiesterase, ENPP1, has been proposed as a mechanism by which highly metastatic tumors evade immune surveillance programs while simultaneously inducing an immunosuppressive profile ^59^. Thus, ENPP1 inhibitors are currently under investigation in preclinical models as well as clinical trials for patients with advanced, unresectable or metastatic tumors.

Conversely, mechanisms by which tumor cells directly modulate DNA sensing and activity of cGAS remain poorly understood. The discovery of c-Src mediated phosphorylation and inhibition of cGAS highlights an important strategy of tumor immune evasion and may provide mechanistic insights into how c-Src inhibition improves T-cell infiltration and immune responses in immunogenic tumors ^67^. Moreover, our data suggests that c-Src inhibitors could be employed as adjuvants or as a rescue strategy to improve tumor immunogenicity during immune checkpoint blockade and/or radiation therapy ^68^. Although not tested here, cGAS tyrosine phosphorylation by other oncogenic SFKs may be a common strategy employed by cancers to promote an immunosuppressive phenotype. Indeed, inhibition of the myeloid-specific SFK hematopoietic cell kinase (HCK) was recently shown to license T-cell mediated immunotherapies in refractory tumors through remodeling of the immunosuppressive TME ^67^.

More generally, it is intriguing to speculate that other growth factor receptor tyrosine kinases and oncogenic signaling pathways may also exhibit inhibitory effects on innate immune sensing pathways in order to promote a non-inflamed TME. In support of this, oncogenic MYC signaling has been shown to induce epigenetic silencing of cGAS-STING signaling in triple-negative breast cancer ^69^. Similarly, the HER2 (ERBB2) pathway disrupts STING signaling through AKT mediated phosphorylation and dysregulation of TBK1, while combination targeted therapy with the tyrosine kinase inhibitor Osimertinib and monoclonal antibodies targeting HER3 was recently shown to induce enhanced tumor cell cGAMP production and paracrine STING activation in tumor-associated macrophages to facilitate tumor clearance^33,70^. Thus, a more detailed understanding of cancer subtype specific strategies of immune evasion will enable the generation of rationally designed combination therapies aimed at restoring tumor immunogenicity.

## Acknowledgements

We would like to acknowledge the Woodward lab for helpful discussions. The Woodward laboratory was supported by National Institutes of Health Grant 1R21AI153820-01. S.A.Z. is supported by the Cora May Poncin fellowship fund. Research reported in this publication was supported by the Office of the Director, National Institutes of Health under Award Number S10OD026741. The content is solely the responsibility of the authors and does not necessarily represent the official views of the National Institutes of Health. The results published here are in part based upon data generated by the TCGA Research Network: https://www.cancer.gov/tcga.

## Author Contributions

Conceived and designed the experiments: W.D., S.A.Z. and J.J.W. Performed the experiments: W.D., S.A.Z., and C.J.H. Analyzed the data: W.D. and S.A.Z. Wrote the paper: W.D., S.A.Z. and J.J.W.

## Quantification and Statistical Analysis

Statistical parameters are reported in the figures and corresponding figure legends. Student t test was used to determine statistical significance * *p* ≤ 0.05; ** *p* ≤ 0.005; ; *** *p* ≤ 0.0005; ns: not significant. The results were expressed as mean ± SD. Value of n is depicted in each figure legend. Luciferase assays were performed using the BioTek synergy HTX plate reader. FRET assays were performed using the CLARIOstar plate reader. qPCR was performed using the CFX Connect Real-Time system (Bio-Rad). Immunoblots were developed using the Odyssey Li-cor (Licor). Phosphor imaging was performed on the Azure Sapphire.

## STAR Methods

### Cells and Viruses

A549 (ATCC), HEK293T (ATCC), and L929-ISRE-luciferase (LUC) ^41^ cells were maintained in Dulbecco’s modified Eagle medium (DMEM; Invitrogen) supplemented with 10% fetal bovine serum (FBS; Genesee). THP-1 (ATCC) cells were grown in RPMI 1640 (Invitrogen) medium supplemented with 10% fetal bovine serum (FBS; Genesee). All cells were maintained with 100 U of penicillin/ml and 100 μg of streptomycin/ml (Invitrogen) at 37 °C under 5% CO_2_. THP-1 cells were differentiated with 150 nM phorbol 12-myristate 13-acetate (PMA) for 24 h followed by removing drug and resting cells in fresh media for 24 h.

### c-Src Inhibitor Treatment

Cells were treated with 30 nM dasatinib (Sigma), 80 nM saracatinib (Selleck), or vehicle for 48 h. Each day, media was replaced with fresh drug. Following drug treatment, experiments were carried out with fresh drug. c-Src inhibition by dasatinib was verified in A549 cells by analyzing c-Src Y416 phosphorylation by Western blot.

### Nucleic Acid Ligand and Plasmid Transfection

Cells were transfected at 70-80% confluency with 10 μg/ml HT-DNA (Sigma), 3 μg/ml ISD45 (Integrated DNA Technologies), 1 μg/ml poly(I:C) (Invivogen), or 2 μg plasmid DNA (6-well plate) using Lipofectamine 2000 (Invitrogen). HT-DNA and ISD45 samples were collected 6 h post-transfection while poly(I:C) samples were collected 16 h post-transfection. Cells transfected with plasmid DNA had fresh media added 6 h post-transfection.

HEK293T cells were transfected at 60-80% confluency with 1 μg (6-well) or 6 μg (10 cm dish) plasmid DNA using 4 μg (6-well) or 24 μg (10 cm dish) PEI (Polysciences). Samples were collected 24 h post-transfection.

### ISRE-Luciferase Assay

HEK293T and A549 cells that had been transfected with HT-DNA, ISD45, or vehicle for 6 h were co-cultured with L929-ISRE-LUC cells for 24 h at 37 °C unless otherwise stated. Cells were then lysed in luciferase lysis buffer (50 mM Tris [pH 7.5], 150 mM NaCl, 1 mM EDTA, 1% NP-40) and cell debris was pelleted by pop-spun centrifugation. The supernatant was transferred to a white-walled 96 well plate and combined with luciferase buffer (20 mM Tricine, 2.67 mM MgSO, 4.7 H2O, 0.1 mM EDTA, 33.3 mM DTT, 530 μM ATP, 270 μM acetyl CoA lithium salt, 470 μM luciferin, 5 mM NaOH, 265 μM magnesium carbonate hydroxide). Luciferase activity was monitored by a BioTek synergy HTX plate reader.

### Cloning and Cell Line Generation

The human c-Src gene was amplified by PCR from the pJP1520-SRC retroviral expression vector (DNASU) and restriction digest cloned into pcDNA3 vector to generate c-Src-HA-pcDNA3. c-Src-HA-pcDNA3 underwent site-directed mutagenesis (SDM) to generate constitutively active (Y530F) and kinase dead (K298M and F408G/Y530F) mutants.

WT and mutant c-Src was further amplified from c-Src-HA-pcDNA3 and restriction digest cloned into pSB-tet-RB ^45^. A549 and THP-1 cells were transfected with WT and mutant HA-c-Src pSB-tet-RB and SB100X (Addgene) followed by blasticidin (Gibco) selection. His-HA-c-Src pSB-tet-RB was generated by SDM and transfected into HEK293T with SB100X followed by blasticidin selection for protein purification.

Human cGAS was amplified by PCR from cGAS pcDNA3 (generously provided by Dr. Genhong Cheng) and restriction digest cloned into pcDNA3 vector to generate FLAG-cGAS-pcDNA3. FLAG-cGAS-pcDNA3 underwent SDM to generate Y248E/F mutants. pSUMO2 human cGAS-FL (generously provided by Dr. Philip Kranzusch) was subjected to SDM to insert a FLAG tag and generate His-FLAG-cGAS pSUMO2 for protein purification.

### Western Blotting

Cell lysates were prepared with lysis buffer (50 mM Tris [pH 7.5], 150 mM NaCl, 1 mM EDTA, 1% NP-40, protease inhibitor cocktail (Thermo)) and quantified by BCA assay (Thermo). Equivalent amounts of each sample were resolved by SDS-PAGE, electrotransferred to nitrocellulose membrane, and blotted for the indicated proteins (Table S1). Primary antibodies were followed by AlexaFluor 680-or -800 conjugated anti-rabbit and anti-mouse secondary antibodies (Li-Cor; 1:10,000) and visualized by Li-Cor Odyssey.

### Nucleic Acid Isolation and Measurement

For analysis of gene expression by RT-qPCR, total RNA was isolated with the Takara NucleoSpin RNA plus kit in accordance with the manufacturer’s instructions. cDNA was synthesized from RNA with random primer (Invitrogen) and Maxima H minus RT (Thermo). qPCR was performed using the PowerUp SYBR Green qPCR kit (Thermo Scientific) with appropriate primers (Table S1).

### CRISPR-Cas9 and pLKO-tet-On-shRNA Cloning, Lentiviral Production, and Infection

Single-guide RNAs (sgRNAs) (sequences in Table S1) were selected by inputting the cGAS gene sequence into the Broad Institute GPP sgRNA CRISPR KO Designer. Two high-score sgRNAs were selected. sgRNA oligos were cloned into the lentiCRISPR V2 (Addgene) lentiviral plasmid according to depositor instructions.

Short-hairpin RNAs (shRNAs) (sequences in Table S1) were selected using the Broad Institute Genetic Perturbation Platform. Two high-score shRNAs were selected for c-Src. shRNA oligos were cloned into the pLKO-Tet-On-shRNA (Addgene) lentiviral plasmid according to depositor instructions.

Lentivirus was prepared in HEK293T cells. Cells were transfected at 50-60% confluency with lentiCRISPR V2, psPAX2 (lentiviral packaging), and pVSV-G (lentiviral envelope) (Addgene) using PEI. 72 h post-transfection the supernatant was collected, mixed with 8 μg/ml of polybrene and 0.5% PEG, and added to HEK293T or A549 cells that were spinfected at 1,000 rpm for 1 h at room temperature. Cells were selected for 2 weeks in media containing 2 μg/ml puromycin (Thermo).

### Knockout Generation

Cells were infected with lentivirus. Following puromycin selection, single cell clones were grown out in 96-well plates. The knockout was verified by transfecting 1 μg/ml poly(I:C) transfected into cells for 16 hr, followed by lysis and analysis of cGAS levels by Western blot. CRISPR-Cas9 induced mutations were identified by isolation of genomic DNA (Promega), PCR of the genomic region where the guide RNA targets, TOPO cloning, and Sanger sequencing of the PCR product.

### NF-kB Activity Luciferase Assay

HEK293T were transfected with 10% pNiFty-LUC (Invivogen), 5% eGFP pcDNA3 (Addgene), FLAG-cGAS pcDNA3, and various c-Src-HA pcDNA3 mutants for 24 h. Luciferase activity was analyzed similarly as the ISRE-luciferase assay. GFP levels were analyzed as a transfection control.

### Co-immunoprecipitations

HEK293T cells were transfected in a 10 cm dish with equal concentrations of EV (pcDNA3), FLAG-cGAS pcDNA3, or c-Src-HA pcDNA3. 24 h post-transfection, cells were collected and lysed in co-IP lysis buffer (50 mM Tris [pH 7.5], 150 mM NaCl, 10% glycerol, 0.5% NP-40) followed by centrifugation. Soluble cell extract was incubated with FLAG M2 magnetic beads (Sigma) or HA magnetic beads (Thermo) for 1 h at 4 °C under rotation. Beads were washed three times for 10 min with co-IP lysis buffer. Resin was eluted with 1X FLAG peptide (Sigma) or HA peptide (Thermo) in co-IP buffer for 1 h at 4 °C. co-IP’d proteins were analyzed by Western blot.

THP-1 WT HA-c-Src pSB-tet-RB cells were treated with 1 μg/ml doxycycline for 48 h and transfected with 1 μg/ml poly(I:C) for 16 h. The co-IP was executed similarly to the HEK293T co-IP and resin was eluted with HA peptide.

4 μg of purified FLAG-cGAS and HA-c-Src were incubated in *in vitro* co-IP buffer (20 mM Tris [pH 7.5], 150 mM NaCl, 3 mM EDTA, 3 mM EGTA, 0.5% NP-40, protease inhibitor cocktail) for 1 h. Proteins were added to FLAG M2 magnetic beads and incubated overnight at 4 °C under rotation. Beads were washed three times for 10 min with *in vitro* co-IP lysis buffer. Resin was eluted with 1X FLAG peptide.

### Purification of FLAG-cGAS, HA-cSrc mutants, BioSTING

Human FLAG-cGAS was purified as previously described ^71^. Briefly, His-FLAG-cGAS pSUMO2 plasmid was transformed into LOBSTR E. coli expression competent cells. Transformed bacteria cultures were inoculated into 1.5 L of LB broth and grown at 37 °C overnight. At OD_600_ 0.5–0.7, protein expression was induced by the addition of 0.5 mM IPTG for 16 h at 16 °C. Bacteria were centrifuged and the cell pellets were resuspended in cGAS lysis buffer (20 mM HEPES [pH 7.5], 400 mM NaCl, 10% glycerol, 30 mM imidazole, 1 mM DTT, 1 mM AEBSF, 0.2 mg/ml lysozyme). Cells were lysed by sonication and clarified lysate was bound to His-Pur Ni-NTA Resin (Thermo). Resin was washed with cGAS lysis buffer, wash buffer (20 mM HEPES [pH 7.5], 1 M NaCl, 10% glycerol, 30 mM imidazole, 1 mM DTT, 0.5 mM AEBSF), lysis buffer, and eluted in cGAS elution buffer (20 mM HEPES [pH 7.5], 400 mM NaCl, 10% glycerol, 300 mM imidazole, 1 mM DTT). Protein was subjected to dialysis with purified human SENP2 SUMO protease in dialysis buffer (20 mM HEPES [pH 7.5], 300 mM NaCl, 1 mM DTT) at 4 °C overnight. FLAG-cGAS was then bound to a heparin column (HiTrap 5 ml Heparin HP) on the AKTA purifier FPLC machine and eluted by buffer B gradient ((buffer A: 20 mM HEPES [pH 7.5), 1 mM DTT) (buffer B: 20 mM HEPES [pH 7.5), 1 M NaCl, 1 mM DTT). Fractions were pooled and concentrated. FLAG-cGAS was further purified by size exclusion chromatography (HiPrep 16/60 Sephacryl S-200) and stored in cGAS storage buffer (20 mM HEPES [pH 7.5], 250 mM KCl, 10% glycerol, 1 mM TCEP). Purity was analyzed by SDS-PAGE followed by Coomassie Brilliant Blue staining. Following concentration, aliquots were flash frozen and stored at −80 °C. Activity was analyzed by *in vitro* FRET assay.

Human WT and mutant His-HA-c-Src pSB-tet-RB HEK293T cells were treated with 2 μg/ml doxycycline for 24 h. Cells were treated with 1 μM dasatinib 1 h prior to lysis. Cells were washed in PBS, lysed in c-Src lysis buffer (20 mM Tris [pH 7.5], 1% Triton X-100, 10% glycerol, 400 mM NaCl, 1 mM EDTA, 20 mM imidazole, 1 mM DTT, 1 mM AEBSF, 10 μM sodium vanadate), and centrifuged to remove cell debris. Supernatant was bound to His-Pur Ni-NTA Resin (Thermo). Resin was washed with 120 bed volumes of lysis buffer and eluted in c-Src elution buffer (20 mM Tris [pH 7.5], 1% Triton X-100, 10% glycerol, 400 mM NaCl, 1 mM EDTA, 300 mM imidazole, 1 mM DTT, 1 mM AEBSF, 10 μM sodium vanadate. Eluted protein was concentrated and subjected to buffer exchange into c-Src storage buffer (50mM Tris [pH 7.5], 150mM NaCI, 1 mM EDTA, 1 mM DTT). Purity was analyzed by Coomassie staining. Following concentration, glycerol was added to a final 50% volume and aliquots were stored at −20°C. Activity was analyzed by auto-phosphorylation via *in vitro* kinase assays followed by probing for c-Src Y416 phosphorylation by Western blot.

BioSTING was purified as previously described ^53^. Briefly, pET15b-BioSTING plasmid was transformed into LOBSTR E. coli and induced similarly to FLAG-cGAS. Following centrifugation, pellet was resuspended in Buffer D (50 mM Tris [pH 7.5], 100 mM NaCl, 20 mM Imidazole, 5 mM β-Mercaptoethanol (BME), 1 mM AEBSF, 0.2 mg/ml lysozyme). Cells were lysed by sonication and clarified lysate was bound to His-Pur Ni-NTA Resin (Thermo). Resin was washed with 120 bed volumes of Buffer D and BioSTING was eluted in Buffer E (50 mM Tris [pH 7.5], 100 mM NaCl, 300 mM Imidazole, 5 mM β-Mercaptoethanol (BME),1 mM AEBSF). Eluted protein was concentrated and subjected to buffer exchange into BioSTING storage buffer (40 mM Tris [pH 7.5], 100 mM NaCl, 30 mM MgCl_2_, 0.5mM TCEP). Purity was analyzed by Coomassie staining. Following concentration, aliquots were flash frozen and stored at −80°C. Activity was analyzed by *in vitro* FRET assay.

### 2’3’ cGAMP enzyme immunoassay (EIA)

HEK293T cells were transfected with FLAG-cGAS pcDNA3 and the indicated c-Src-HA pcDNA3 mutants using PEI for 24 h. Cells were washed with cold PBS and lysates were prepared in accordance with the manufacturer’s instructions (Arbor Assays).

### *In vitro* kinase assay

WT and mutant HA-c-Src was incubated with FLAG-cGAS in kinase buffer (5 mM MOPS [pH 7.2], 2.5 mM ®-glycerol-phosphate, 4 mM MgCl_2_, 2.5 mM MnCl_2_, 1 mM EGTA, 0.4 mM EDTA, 10 mM DTT, 50 μM ATP, 4 μCi ©-32P ATP) at 30 °C for 30 min. Non-radioactive assays were conducted with 250 μM ATP. If end-point assay, reaction was stopped by adding 4X SDS-PAGE loading dye at 95 °C for 5 min following by SDS-PAGE analysis. Radioactive SDS-PAGE gels were imaged on an Azure Sapphire imager.

### FRET assays

Cellular: HEK293T cells were transfected with FLAG-cGAS pcDNA3 and indicated c-Src-HA pcDNA3 mutants using PEI for 24 h. Cells were lysed in lysis buffer (50 mM Tris [pH 7.5], 150 mM NaCl, 1mM EDTA, 1% NP-40, protease inhibitor cocktail (Thermo), centrifuged to remove cell debris, and supernatants were incubated with 1 μM of purified BioSTING in BioSTING activity buffer (50 mM Tris [pH 7.5], 35 mM KCl, 5 mM Mg(OAc)_2_). Enzyme assays were allowed to proceed for 2 h at 37 °C and FRET fluorescence was monitored using a FRET-capable plate reader (CLARIOstar) with the following parameters: 458 nm excitation, 490 nm and 600 nm emission.

*In vitro*: 30 nM FLAG-cGAS was phosphorylated by WT HA-c-Src using the above *in vitro* kinase assay. The reaction was combined with 1 μM of purified BioSTING, 0.5 ng ISD45, 25 μM ATP, 25 μM GTP in BioSTING activity buffer (50 mM Tris [pH 7.5], 35 mM KCl, 5 mM Mg(OAc)_2_).

### FLAG-cGAS ChIP

HEK293T cells were transfected with FLAG-cGAS pcDNA3, EV pcDNA3, and c-Src-HA pcDNA3 using PEI for 24 h. Cells were crosslinked in 1% formaldehyde (VWR) for 10 min and quenched with glycine prior to snap-freezing. Cell pellets were lysed in SDS lysis buffer (0.8% SDS, 10 mM EDTA, 50 mM Tris [pH 8.1], protease inhibitor cocktail), sonicated, and centrifuged to remove cell debris. Supernatant was diluted 3-fold in dilution buffer (0.003% SDS, 0.4 mM EDTA, 5.6 mM Tris [pH 8.1], 3.3% Triton X-100, 501 mM NaCl, protease inhibitor cocktail), and incubated with FLAG M2 magnetic beads overnight at 4 °C under rotation. Beads were washed two times in low salt wash buffer (0.1% SDS, 2 mM EDTA, 20 mM Tris [pH 8.1], 1% Triton X-100, 150 mM NaCl), two times in high salt wash buffer (0.1% SDS, 2 mM EDTA, 20 mM Tris [pH 8.1], 1% Triton X-100, 500 mM NaCl), two times in LiCl wash buffer (0.25 M LiCl, 1 mM EDTA, 10 mM Tris [pH 8.1], 1% NP-40, 1% sodium deoxycholic acid), and two times in TE buffer (1 mM EDTA, 10 mM Tris [pH 8.1]). Input and beads were resuspended in elution buffer (1% SDS, 150 mM NaCl) supplemented with 150 μg/ml 1X FLAG peptide and crosslinks were reversed by incubating at 65 °C overnight. Following bead removal by magnet, DNA was isolated from 90% of the supernatant by adding 50 μg/ml proteinase K (Thermo) and RNase A (NEB) at 65 °C for 2 h followed by PCR clean-up. Bound pcDNA3 was analyzed by qPCR. The remaining 10% was treated with DNase (Invitrogen) and RNase A at 37 °C and analyzed by Western blot.

### Biotin-ISD45 Annealing and Immunoprecipitation

Biotinylated forward and non-modified reverse ISD45 oligos (Integrated DNA Technologies) were resuspended at 1 mg/ml in annealing buffer (5 mM Tris [pH 7.5], 25 mM NaCl) and annealed by incubating oligos together at 95 °C for 5 min then cooled to 25 °C at 0.1 °C/sec.

HEK293T cells were transfected with FLAG-cGAS pcDNA3, EV pcDNA3, and c-Src-HA pcDNA3 for 24 h. 4 μg/ml annealed ISD45 was transfected into cells using PEI. 2 h later, cells were washed in PBS, lysed in biotin-ISD45 lysis buffer (20 mM Tris [pH 7.5], 100 mM NaCl, 3mM EDTA, 3mM EGTA, 0.5% NP-40, 0.1% BSA, protease inhibitor cocktail) followed by centrifugation. Cell lysates were incubated with streptavidin magnetic beads (NEB) for 1 h at 4 °C under rotation. Beads were washed five times in lysis buffer and protein was eluted off resin using 1X SDS-PAGE loading dye and heating at 95 °C for 7 min.

### TCGA Correlation Heat Maps

Tumor gene expression data from lung adenocarcinoma (LUAD), melanoma (SKCM), and bladder cancer (BLCA) patients were analyzed from the publicly available The Cancer Genome Atlas (TCGA) database using the UCSC Xena platform (https://xenabrowser.net) ^38^. RNA expression levels of c-Src, IFN-®, IFIT2, PKR, IL6, and FUS were analyzed. Expression range and corresponding colors are included for each gene representing log2(fpkm-uq+1). Pearson’s rho correlation value and p values are listed in Figures 1 and S1.

### Kaplan-Meier Survival Curves

Kaplan–Meier survival curves were generated using publicly available human lung cancer patient microarray datasets (kmplot.com/analysis) ^72^. Patients were divided by cGAS (MB21D1) and c-Src (SRC) expression values. Expression cutoff was dictated by auto select best cutoff in program. FDR, HR, and log-rank *P*-values are listed in the figure and figure legend.

## Supplemental Figure Titles and Legends

**Figure S1: p values from heat maps, and oncogenic B-Raf and K-Ras expression does not correlate with reduced innate immune gene expression in melanoma and lung adenocarcinoma, respectively. (A)** p values for gene expression heat maps in various cancer. **(B)** Heat map of TCGA data of B-Raf, IFN ®, IFIT2, PKR, IL6, and FUS expression levels in melanoma patients. Data was analyzed through the UCSC Xena program. Expression range and corresponding colors are included for each gene representing log2(fpkm-uq+1). **(C)** Heat map of TCGA data for K-Ras and immune gene expression in lung adenocarcinoma patients similar to (B).

**Figure S2: Cyclic dinucleotides including 2’3’ cGAMP are transferred from transfected cells into L929-ISRE-LUC reporter cells by co-culturing via gap junctions. (A)** Co-culture of L929-ISRE-LUC cells with HEK293T cells transfected with pcDNA3 empty vector (EV) or CDN cyclases. **(B)** L929-ISRE-LUC cells were mock treated, mock transfected with lipofectamine 2000, or transfected with 50 μM 3’3’ cGAMP, 50 μM 2’3’ cGAMP, or 1 μg ISD-100mer. Luciferase activity was quantified 18 h later. **(C)** Co-culture of L929-ISRE-LUC cells with HEK293T cells transfected with decreasing concentrations of cGAS-pcDNA3 (1000, 100, 10, 0 ng) and 3 μg of either wildtype (WT) or catalytically inactive (H17A) Poxin. **(D)** Co-culture of L929-ISRE-LUC cells with HEK293T cells transfected with cGAS-pcDNA3 (1000 or 0 ng) in the presence or absence of the gap junction inhibitor meclofenamic acid (MFA) (100 uM). *n* = 4 biological replicates.

**Figure S3: c-Src, cGAS, and STING are expressed at different levels in various cell lines and impacts whether c-Src inhibition or overexpression regulates DNA sensing. (A)** Western blot of HEK293T, A549, and THP-1 cells treated with 1 μg/ml poly(I:C) for 16 h. ®-actin was used as a loading control. **(B)** Western blot of A549 cells treated with 30 nM dasatinib or vehicle for 48 h. p Y416 is a marker for c-Src autophosphorylation and activation. ®-actin was used as a loading control. **(C)** Co-culture of L929-ISRE-LUC cells with A549 cells treated with vehicle, dasatinib (30nM), or saracatinib (80 nM) for 48 h followed by transfection with HT-DNA for 6 h. **(D)** RT-qPCR analysis of IFIT2 levels from A549 cells transfected with EV or HA-STING pcDNA3 and treated with vehicle or dasatinib for 48 h followed by transfection with indicated DNA ligands for 6 h. All samples were normalized to 18S rRNA. **(E)** Western blot of A549 WT and mutant HA-c-Src-tet-RB SB cells treated with 1 μg/ml doxycycline (DOX) for 48 h. ®-actin was used as a loading control. **(F)** Co-culture of L929-ISRE-LUC cells with cells in (E) followed by transfection with indicated DNA ligands for 6 h. **(G)** RT-qPCR analysis of IFN ® levels from THP-1 cells treated with vehicle or dasatinib for 48 h followed by transfection with indicated DNA ligands for 6 h. All samples were normalized to 18S rRNA. *n*= 4 biological replicates for luciferase assay, and *n*= 3 biological replicates and representative qPCR with technical triplicates is shown.

**Figure S4: c-Src inhibits cGAS-dependent NF-**|**B activity and c-Src depletion enhances a DNA-dependent IFN response. (A)** NF-|B luciferase assay from HEK293T cells transfected with pNiFty-LUC, cGAS pcDNA3, and EV or c-Src-HA pcDNA3 WT or mutant pcDNA3 for 24 h. NF-|B activity was analyzed by luciferase activity and normalized to transfected eGFP plasmid. **(B)** RT-qPCR analysis of c-Src expression in HEK293T cells transduced with DOX-inducible non-target (CON) or independent c-Src-targeting shRNAs. DOX was added for 72 h. c-Src levels in CON were set to 1. All samples were normalized to 18S rRNA. **(C)** Co-culture of L929-ISRE-LUC cells with cells in (B) transfected with cGAS pcDNA3 for 24 h followed by transfection of ISD45 or 2’3’ cGAMP for 6 h. Cells not transfected with cGAS were used as a negative control (Negative). *n*= 4 biological replicates for luciferase assays, and *n*= 3 biological replicates and representative qPCR with technical triplicates is shown.

**Figure S5: Human FLAG-cGAS and HA-c-Src purification and activity validation. (A)** Coomassie brilliant blue stain of purified FLAG-cGAS after SEC and concentration. **(B)** BioSTING FRET assay using various concentration of purified FLAG-cGAS for 2 h. **(C)** Coomassie brilliant blue stain of purified WT and mutant HA-c-Src after concentration. Arrow depicts not-specific band. **(D)** Western blot of purified WT and mutant HA-c-Src *in vitro* kinase assays using cold ATP. c-Src Y416 designates autophosphorylation and c-Src kinase activity. *n*= 2 biological replicates.

**Table S1: Primer and Oligo Sequences**

## References

1. Tang, D., Kang, R., Coyne, C.B., Zeh, H.J., and Lotze, M.T. (2012). PAMPs and DAMPs: signal 0s that spur autophagy and immunity. Immunol. Rev. 249, 158–175. 10.1111/j.1600-065X.2012.01146.x.

2. Sun, L., Wu, J., Du, F., Chen, X., and Chen, Z.J. (2013). Cyclic GMP-AMP synthase is a cytosolic DNA sensor that activates the type I interferon pathway. Science 339, 786–791. 10.1126/science.1232458.

3. Wu, J., Sun, L., Chen, X., Du, F., Shi, H., Chen, C., and Chen, Z.J. (2013). Cyclic GMP-AMP is an endogenous second messenger in innate immune signaling by cytosolic DNA. Science 339, 826–830. 10.1126/science.1229963.

4. Takaoka, A., Wang, Z., Choi, M.K., Yanai, H., Negishi, H., Ban, T., Lu, Y., Miyagishi, M., Kodama, T., Honda, K., et al. (2007). DAI (DLM-1/ZBP1) is a cytosolic DNA sensor and an activator of innate immune response. Nature 448, 501–505. 10.1038/nature06013.

5. Bürckstümmer, T., Baumann, C., Blüml, S., Dixit, E., Dürnberger, G., Jahn, H., Planyavsky, M., Bilban, M., Colinge, J., Bennett, K.L., et al. (2009). An orthogonal proteomic-genomic screen identifies AIM2 as a cytoplasmic DNA sensor for the inflammasome. Nat. Immunol. 10, 266–272. 10.1038/ni.1702.

6. Ferguson, B.J., Mansur, D.S., Peters, N.E., Ren, H., and Smith, G.L. (2012). DNA-PK is a DNA sensor for IRF-3-dependent innate immunity. eLife 1, e00047. 10.7554/eLife.00047.

7. Han, F., Guo, H., Wang, L., Zhang, Y., Sun, L., Dai, C., and Wu, X. (2021). The cGAS-STING signaling pathway contributes to the inflammatory response and autophagy in Aspergillus fumigatus keratitis. Exp. Eye Res. 202, 108366. 10.1016/j.exer.2020.108366.

8. Hopfner, K.-P., and Hornung, V. (2020). Molecular mechanisms and cellular functions of cGAS–STING signalling. Nat. Rev. Mol. Cell Biol. 21, 501–521. 10.1038/s41580-020-0244-x.

9. Ablasser, A., Goldeck, M., Cavlar, T., Deimling, T., Witte, G., Röhl, I., Hopfner, K.-P., Ludwig, J., and Hornung, V. (2013). cGAS produces a 2′-5′-linked cyclic dinucleotide second messenger that activates STING. Nature 498, 380–384. 10.1038/nature12306.

10. Zhang, X., Shi, H., Wu, J., Zhang, X., Sun, L., Chen, C., and Chen, Z.J. (2013). Cyclic GMP-AMP containing mixed phosphodiester linkages is an endogenous high-affinity ligand for STING. Mol. Cell 51, 226–235. 10.1016/j.molcel.2013.05.022.

11. Diner, E.J., Burdette, D.L., Wilson, S.C., Monroe, K.M., Kellenberger, C.A., Hyodo, M., Hayakawa, Y., Hammond, M.C., and Vance, R.E. (2013). The innate immune DNA sensor cGAS produces a noncanonical cyclic dinucleotide that activates human STING. Cell Rep. 3, 1355–1361. 10.1016/j.celrep.2013.05.009.

12. Gao, P., Ascano, M., Wu, Y., Barchet, W., Gaffney, B.L., Zillinger, T., Serganov, A.A., Liu, Y., Jones, R.A., Hartmann, G., et al. (2013). Cyclic [G(2’,5’)pA(3’,5’)p] is the metazoan second messenger produced by DNA-activated cyclic GMP-AMP synthase. Cell 153, 1094–1107. 10.1016/j.cell.2013.04.046.

13. Ishikawa, H., and Barber, G.N. (2008). STING is an endoplasmic reticulum adaptor that facilitates innate immune signalling. Nature 455, 674–678. 10.1038/nature07317.

14. Ablasser, A., Hemmerling, I., Schmid-Burgk, J.L., Behrendt, R., Roers, A., and Hornung, V. (2014). TREX1 Deficiency Triggers Cell-Autonomous Immunity in a cGAS-Dependent Manner. J. Immunol. 192, 5993–5997. 10.4049/jimmunol.1400737.

15. Ahn, J., Ruiz, P., and Barber, G.N. (2014). Intrinsic Self-DNA Triggers Inflammatory Disease Dependent on STING. J. Immunol. 193, 4634–4642. 10.4049/jimmunol.1401337.

16. Gray, E.E., Treuting, P.M., Woodward, J.J., and Stetson, D.B. (2015). Cutting Edge: cGAS Is Required for Lethal Autoimmune Disease in the Trex1-Deficient Mouse Model of Aicardi–Goutières Syndrome. J. Immunol. 195, 1939–1943. 10.4049/jimmunol.1500969.

17. Gao, D., Li, T., Li, X.-D., Chen, X., Li, Q.-Z., Wight-Carter, M., and Chen, Z.J. (2015). Activation of cyclic GMP-AMP synthase by self-DNA causes autoimmune diseases. Proc. Natl. Acad. Sci. 112, E5699–E5705. 10.1073/pnas.1516465112.

18. Mackenzie, K.J., Carroll, P., Lettice, L., Tarnauskaitė, Ž., Reddy, K., Dix, F., Revuelta, A., Abbondati, E., Rigby, R.E., Rabe, B., et al. (2016). Ribonuclease H2 mutations induce a cGAS/STING-dependent innate immune response. EMBO J. 35, 831–844. 10.15252/embj.201593339.

19. Liang, Q., Seo, G.J., Choi, Y.J., Kwak, M.-J., Ge, J., Rodgers, M.A., Shi, M., Leslie, B.J., Hopfner, K.-P., Ha, T., et al. (2014). Crosstalk between the cGAS DNA Sensor and Beclin-1 Autophagy Protein Shapes Innate Antimicrobial Immune Responses. Cell Host Microbe 15, 228–238. 10.1016/j.chom.2014.01.009.

20. Hu, M.-M., Yang, Q., Xie, X.-Q., Liao, C.-Y., Lin, H., Liu, T.-T., Yin, L., and Shu, H.-B. (2016). Sumoylation Promotes the Stability of the DNA Sensor cGAS and the Adaptor STING to Regulate the Kinetics of Response to DNA Virus. Immunity 45, 555–569. 10.1016/j.immuni.2016.08.014.

21. Seo, G.J., Yang, A., Tan, B., Kim, S., Liang, Q., Choi, Y., Yuan, W., Feng, P., Park, H.-S., and Jung, J.U. (2015). Akt Kinase-Mediated Checkpoint of cGAS DNA Sensing Pathway. Cell Rep. 13, 440–449. 10.1016/j.celrep.2015.09.007.

22. Zhong, L., Hu, M.-M., Bian, L.-J., Liu, Y., Chen, Q., and Shu, H.-B. (2020). Phosphorylation of cGAS by CDK1 impairs self-DNA sensing in mitosis. Cell Discov. 6, 1–12. 10.1038/s41421-020-0162-2.

23. Li, T., Huang, T., Du, M., Chen, X., Du, F., Ren, J., and Chen, Z.J. (2021). Phosphorylation and chromatin tethering prevent cGAS activation during mitosis. Science 371, eabc5386. 10.1126/science.abc5386.

24. Kwon, J., and Bakhoum, S.F. (2020). The Cytosolic DNA-Sensing cGAS–STING Pathway in Cancer. Cancer Discov. 10, 26–39. 10.1158/2159-8290.CD-19-0761.

25. Yang, H., Wang, H., Ren, J., Chen, Q., and Chen, Z.J. (2017). cGAS is essential for cellular senescence. Proc. Natl. Acad. Sci. U. S. A. 114, E4612–E4620. 10.1073/pnas.1705499114.

26. Tumor-Derived cGAMP Triggers a STING-Mediated Interferon Response in Non-tumor Cells to Activate the NK Cell Response (2018). Immunity 49, 754-763.e4. 10.1016/j.immuni.2018.09.016.

27. Garland, K.M., Rosch, J.C., Carson, C.S., Wang-Bishop, L., Hanna, A., Sevimli, S., Van Kaer, C., Balko, J.M., Ascano, M., and Wilson, J.T. (2021). Pharmacological Activation of cGAS for Cancer Immunotherapy. Front. Immunol. 12, 753472. 10.3389/fimmu.2021.753472.

28. Corrales, L., Glickman, L.H., McWhirter, S.M., Kanne, D.B., Sivick, K.E., Katibah, G.E., Woo, S.-R., Lemmens, E., Banda, T., Leong, J.J., et al. (2015). Direct Activation of STING in the Tumor Microenvironment Leads to Potent and Systemic Tumor Regression and Immunity. Cell Rep. 11, 1018–1030. 10.1016/j.celrep.2015.04.031.

29. Wang, H., Hu, S., Chen, X., Shi, H., Chen, C., Sun, L., and Chen, Z.J. (2017). cGAS is essential for the antitumor effect of immune checkpoint blockade. Proc. Natl. Acad. Sci. U. S. A. 114, 1637–1642. 10.1073/pnas.1621363114.

30. Flood, B.A., Higgs, E.F., Li, S., Luke, J.J., and Gajewski, T.F. (2019). STING pathway agonism as a cancer therapeutic. Immunol. Rev. 290, 24–38. 10.1111/imr.12765.

31. Konno, H., Yamauchi, S., Berglund, A., Putney, R.M., Mulé, J.J., and Barber, G.N. (2018). Suppression of STING Signaling through Epigenetic Silencing and Missense Mutation Impedes DNA-Damage Mediated Cytokine Production. Oncogene 37, 2037–2051. 10.1038/s41388-017-0120-0.

32. Wu, M.-Z., Cheng, W.-C., Chen, S.-F., Nieh, S., O’Connor, C., Liu, C.-L., Tsai, W.-W., Wu, C.-J., Martin, L., Lin, Y.-S., et al. (2017). miR-25/93 mediates hypoxia-induced immunosuppression by repressing cGAS. Nat. Cell Biol. 19, 1286–1296. 10.1038/ncb3615.

33. Wu, S., Zhang, Q., Zhang, F., Meng, F., Liu, S., Zhou, R., Wu, Q., Li, X., Shen, L., Huang, J., et al. (2019). HER2 recruits AKT1 to disrupt STING signalling and suppress antiviral defence and antitumour immunity. Nat. Cell Biol. 21, 1027–1040. 10.1038/s41556-019-0352-z.

34. Ishizawar, R., and Parsons, S.J. (2004). c-Src and cooperating partners in human cancer. Cancer Cell 6, 209–214. 10.1016/j.ccr.2004.09.001.

35. Irby, R.B., and Yeatman, T.J. (2000). Role of Src expression and activation in human cancer. Oncogene 19, 5636–5642. 10.1038/sj.onc.1203912.

36. Belli, S., Esposito, D., Servetto, A., Pesapane, A., Formisano, L., and Bianco, R. (2020). c-Src and EGFR Inhibition in Molecular Cancer Therapy: What Else Can We Improve? Cancers 12, 1489. 10.3390/cancers12061489.

37. Uhlen, M., Zhang, C., Lee, S., Sjöstedt, E., Fagerberg, L., Bidkhori, G., Benfeitas, R., Arif, M., Liu, Z., Edfors, F., et al. (2017). A pathology atlas of the human cancer transcriptome. Science 357, eaan2507. 10.1126/science.aan2507.

38. Goldman, M.J., Craft, B., Hastie, M., Repecka, K., McDade, F., Kamath, A., Banerjee, A., Luo, Y., Rogers, D., Brooks, A.N., et al. (2020). Visualizing and interpreting cancer genomics data via the Xena platform. Nat. Biotechnol. 38, 675–678. 10.1038/s41587-020-0546-8.

39. Ascierto, P.A., Kirkwood, J.M., Grob, J.-J., Simeone, E., Grimaldi, A.M., Maio, M., Palmieri, G., Testori, A., Marincola, F.M., and Mozzillo, N. (2012). The role of BRAF V600 mutation in melanoma. J. Transl. Med. 10, 85. 10.1186/1479-5876-10-85.

40. Yang, H., Liang, S.-Q., Schmid, R.A., and Peng, R.-W. (2019). New Horizons in KRAS-Mutant Lung Cancer: Dawn After Darkness. Front. Oncol. 9, 953. 10.3389/fonc.2019.00953.

41. Woodward, J.J., Iavarone, A.T., and Portnoy, D.A. (2010). c-di-AMP secreted by intracellular Listeria monocytogenes activates a host type I interferon response. Science 328, 1703–1705. 10.1126/science.1189801.

42. Ablasser, A., Schmid-Burgk, J.L., Hemmerling, I., Horvath, G.L., Schmidt, T., Latz, E., and Hornung, V. (2013). Cell intrinsic immunity spreads to bystander cells via the intercellular transfer of cGAMP. Nature 503, 530–534. 10.1038/nature12640.

43. Eaglesham, J.B., Pan, Y., Kupper, T.S., and Kranzusch, P.J. (2019). Viral and metazoan poxins are cGAMP-specific nucleases that restrict cGAS-STING signalling. Nature 566, 259–263. 10.1038/s41586-019-0928-6.

44. Kmiecik, T.E., Johnson, P.J., and Shalloway, D. (1988). Regulation by the autophosphorylation site in overexpressed pp60c-src. Mol. Cell. Biol. 8, 4541–4546. 10.1128/mcb.8.10.4541-4546.1988.

45. Kowarz, E., Löscher, D., and Marschalek, R. (2015). Optimized Sleeping Beauty transposons rapidly generate stable transgenic cell lines. Biotechnol. J. 10, 647–653. 10.1002/biot.201400821.

46. Agius, M.P., Ko, K.S., Johnson, T.K., Kwarcinski, F.E., Phadke, S., Lachacz, E.J., and Soellner, M.B. (2019). Selective Proteolysis to Study the Global Conformation and Regulatory Mechanisms of c-Src Kinase. ACS Chem. Biol. 14, 1556–1563. 10.1021/acschembio.9b00306.

47. Cartwright, C.A., Eckhart, W., Simon, S., and Kaplan, P.L. (1987). Cell transformation by pp60c-src mutated in the carboxy-terminal regulatory domain. Cell 49, 83–91. 10.1016/0092-8674(87)90758-6.

48. Kmiecik, T.E., and Shalloway, D. (1987). Activation and suppression of pp60c-src transforming ability by mutation of its primary sites of tyrosine phosphorylation. Cell 49, 65–73. 10.1016/0092-8674(87)90756-2.

49. Piwnica-Worms, H., Saunders, K.B., Roberts, T.M., Smith, A.E., and Cheng, S.H. (1987). Tyrosine phosphorylation regulates the biochemical and biological properties of pp60c-src. Cell 49, 75–82. 10.1016/0092-8674(87)90757-4.

50. Snyder, M.A., Bishop, J.M., McGrath, J.P., and Levinson, A.D. (1985). A mutation at the ATP-binding site of pp60v-src abolishes kinase activity, transformation, and tumorigenicity. Mol. Cell. Biol. 5, 1772–1779. 10.1128/mcb.5.7.1772-1779.1985.

51. Parsons, S.J., and Parsons, J.T. (2004). Src family kinases, key regulators of signal transduction. Oncogene 23, 7906–7909. 10.1038/sj.onc.1208160.

52. Rehwinkel, J., and Gack, M.U. (2020). RIG-I-like receptors: their regulation and roles in RNA sensing. Nat. Rev. Immunol. 20, 537–551. 10.1038/s41577-020-0288-3.

53. Pollock, A.J., Zaver, S.A., and Woodward, J.J. (2020). A STING-based biosensor affords broad cyclic dinucleotide detection within single living eukaryotic cells. Nat. Commun. 11, 3533. 10.1038/s41467-020-17228-y.

54. Guey, B., Wischnewski, M., Decout, A., Makasheva, K., Kaynak, M., Sakar, M.S., Fierz, B., and Ablasser, A. (2020). BAF restricts cGAS on nuclear DNA to prevent innate immune activation. Science 369, 823–828. 10.1126/science.aaw6421.

55. Hornbeck, P.V., Zhang, B., Murray, B., Kornhauser, J.M., Latham, V., and Skrzypek, E. (2015). PhosphoSitePlus, 2014: mutations, PTMs and recalibrations. Nucleic Acids Res. 43, D512–520. 10.1093/nar/gku1267.

56. Liu, H., Zhang, H., Wu, X., Ma, D., Wu, J., Wang, L., Jiang, Y., Fei, Y., Zhu, C., Tan, R., et al. (2018). Nuclear cGAS suppresses DNA repair and promotes tumorigenesis. Nature 563, 131–136. 10.1038/s41586-018-0629-6.

57. Jiang, H., Xue, X., Panda, S., Kawale, A., Hooy, R.M., Liang, F., Sohn, J., Sung, P., and Gekara, N.O. (2019). Chromatin-bound cGAS is an inhibitor of DNA repair and hence accelerates genome destabilization and cell death. EMBO J. 38, e102718. 10.15252/embj.2019102718.

58. Zhang, X., Wu, J., Du, F., Xu, H., Sun, L., Chen, Z., Brautigam, C.A., Zhang, X., and Chen, Z.J. (2014). The cytosolic DNA sensor cGAS forms an oligomeric complex with DNA and undergoes switch-like conformational changes in the activation loop. Cell Rep. 6, 421–430. 10.1016/j.celrep.2014.01.003.

59. Li, J., Duran, M.A., Dhanota, N., Chatila, W.K., Bettigole, S.E., Kwon, J., Sriram, R.K., Humphries, M.P., Salto-Tellez, M., James, J.A., et al. (2021). Metastasis and immune evasion from extracellular cGAMP hydrolysis. Cancer Discov. 11, 1212–1227. 10.1158/2159-8290.CD-20-0387.

60. Mekers, V.E., Kho, V.M., Ansems, M., and Adema, G.J. (2022). cGAS/cGAMP/STING signal propagation in the tumor microenvironment: Key role for myeloid cells in antitumor immunity. Radiother. Oncol. J. Eur. Soc. Ther. Radiol. Oncol. 174, 158–167. 10.1016/j.radonc.2022.07.014.

61. Campisi, M., Sundararaman, S.K., Shelton, S.E., Knelson, E.H., Mahadevan, N.R., Yoshida, R., Tani, T., Ivanova, E., Cañadas, I., Osaki, T., et al. (2020). Tumor-Derived cGAMP Regulates Activation of the Vasculature. Front. Immunol. 11, 2090. 10.3389/fimmu.2020.02090.

62. Huang, K.-C., Chanda, D., McGrath, S., Dixit, V., Zhang, C., Wu, J., Tendyke, K., Yao, H., Hukkanen, R., Taylor, N., et al. (2022). Pharmacologic Activation of STING in the Bladder Induces Potent Antitumor Immunity in Non-Muscle Invasive Murine Bladder Cancer. Mol. Cancer Ther. 21, 914–924. 10.1158/1535-7163.MCT-21-0780.

63. Lombardo, K.A., Obradovic, A., Singh, A.K., Liu, J.L., Joice, G., Kates, M., Bishai, W., McConkey, D., Chaux, A., Eich, M.-L., et al. (2022). BCG invokes superior STING-mediated innate immune response over radiotherapy in a carcinogen murine model of urothelial cancer. J. Pathol. 256, 223–234. 10.1002/path.5830.

64. Singh, A.K., Praharaj, M., Lombardo, K.A., Yoshida, T., Matoso, A., Baras, A.S., Zhao, L., Srikrishna, G., Huang, J., Prasad, P., et al. (2022). Re-engineered BCG overexpressing cyclic di-AMP augments trained immunity and exhibits improved efficacy against bladder cancer. Nat. Commun. 13, 878. 10.1038/s41467-022-28509-z.

65. Li, S., Mirlekar, B., Johnson, B.M., Brickey, W.J., Wrobel, J.A., Yang, N., Song, D., Entwistle, S., Tan, X., Deng, M., et al. (2022). STING-induced regulatory B cells compromise NK function in cancer immunity. Nature. 10.1038/s41586-022-05254-3.

66. Falahat, R., Berglund, A., Putney, R.M., Perez-Villarroel, P., Aoyama, S., Pilon-Thomas, S., Barber, G.N., and Mulé, J.J. (2021). Epigenetic reprogramming of tumor cell–intrinsic STING function sculpts antigenicity and T cell recognition of melanoma. Proc. Natl. Acad. Sci. U. S. A. 118, e2013598118. 10.1073/pnas.2013598118.

67. Poh, A.R., Love, C.G., Chisanga, D., Steer, J.H., Baloyan, D., Chopin, M., Nutt, S., Rautela, J., Huntington, N.D., Etemadi, N., et al. Therapeutic inhibition of the SRC-kinase HCK facilitates T cell tumor infiltration and improves response to immunotherapy. Sci. Adv. 8, eabl7882. 10.1126/sciadv.abl7882.

68. Redin, E., Garmendia, I., Lozano, T., Serrano, D., Senent, Y., Redrado, M., Villalba, M., De Andrea, C.E., Exposito, F., Ajona, D., et al. (2021). SRC family kinase (SFK) inhibitor dasatinib improves the antitumor activity of anti-PD-1 in NSCLC models by inhibiting Treg cell conversion and proliferation. J. Immunother. Cancer 9, e001496. 10.1136/jitc-2020-001496.

69. Wu, S.-Y., Xiao, Y., Wei, J.-L., Xu, X.-E., Jin, X., Hu, X., Li, D.-Q., Jiang, Y.-Z., and Shao, Z.-M. (2021). MYC suppresses STING-dependent innate immunity by transcriptionally upregulating DNMT1 in triple-negative breast cancer. J. Immunother. Cancer 9, e002528. 10.1136/jitc-2021-002528.

70. Vicencio, J.M., Evans, R., Green, R., An, Z., Deng, J., Treacy, C., Mustapha, R., Monypenny, J., Costoya, C., Lawler, K., et al. (2022). Osimertinib and anti-HER3 combination therapy engages immune dependent tumor toxicity via STING activation in trans. Cell Death Dis. 13, 274. 10.1038/s41419-022-04701-3.

71. Zhou, W., Whiteley, A.T., de Oliveira Mann, C.C., Morehouse, B.R., Nowak, R.P., Fischer, E.S., Gray, N.S., Mekalanos, J.J., and Kranzusch, P.J. (2018). Structure of the Human cGAS-DNA Complex Reveals Enhanced Control of Immune Surveillance. Cell 174, 300-311.e11. 10.1016/j.cell.2018.06.026.

72. Győrffy, B., Surowiak, P., Budczies, J., and Lánczky, A. (2013). Online survival analysis software to assess the prognostic value of biomarkers using transcriptomic data in non-small-cell lung cancer. PloS One 8, e82241. 10.1371/journal.pone.0082241.

