## Supplemental Figures for "The proto-oncogene c-Src phosphorylates cGAS to reduce DNA binding and activation"

A)

**p Values**

| | | IFN B | IFIT2 | PKR | IL6 | FUS |
|---|---|---|---|---|---|---|
| c-Src | Melanoma | 0.002511 | 0.000566 | 6.35E-08 | 0.006979 | 2.45E-08 |
| | Bladder | 8.91E-05 | 1.36E-09 | 7.55E-05 | 3.23E-16 | 3.56E-05 |
| | Lung | 0.04415 | 0.009475 | 0.01661 | 1.98E-09 | 0.001755 |
| B-Raf | Melanoma | 0.321 | 6.66E-07 | 4.14E-18 | 0.3854 | 4.7E-19 |
| K-Ras | Lung | 0.5518 | 0.000189 | 3.53E-13 | 0.9546 | 0.3285 |

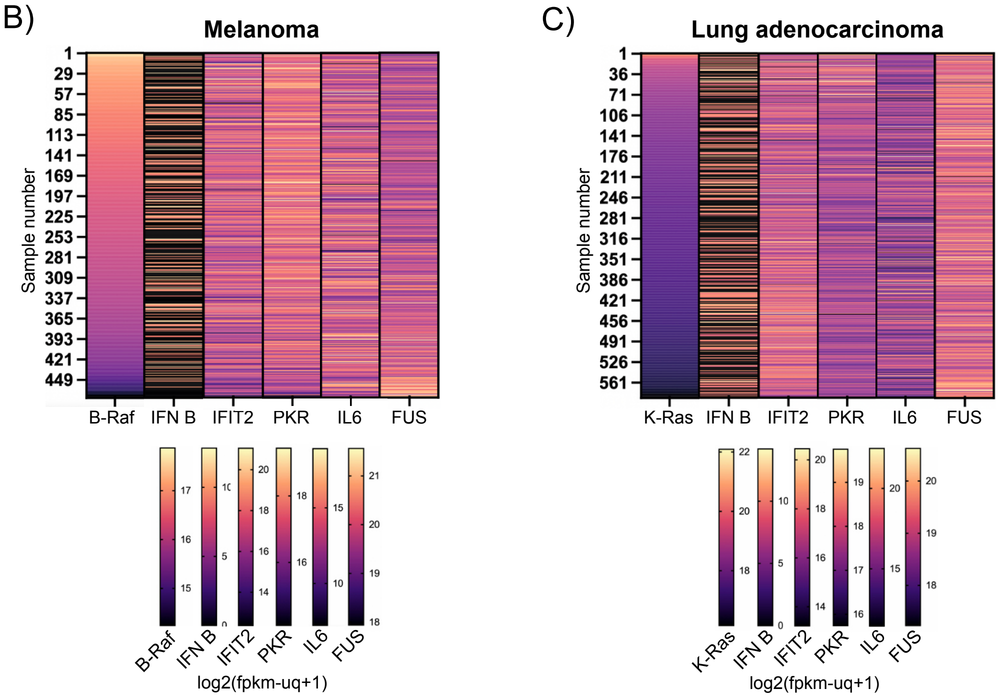

**Figure S1: p values from heat maps, and oncogenic B-Raf and K-Ras expression does not correlate with reduced innate immune gene expression in melanoma and lung adenocarcinoma, respectively. (A)** p values for gene expression heat maps in various cancer. **(B)** Heat map of TCGA data of B-Raf, IFN b , IFIT2, PKR, IL6, and FUS expression levels in melanoma patients. Data was analyzed through the UCSC Xena program. Expression range and corresponding colors are included for each gene representing log2(fpkm-uq+1). **(C)** Heat map of TCGA data for K-Ras and immune gene expression in lung adenocarcinoma patients similar to (B).

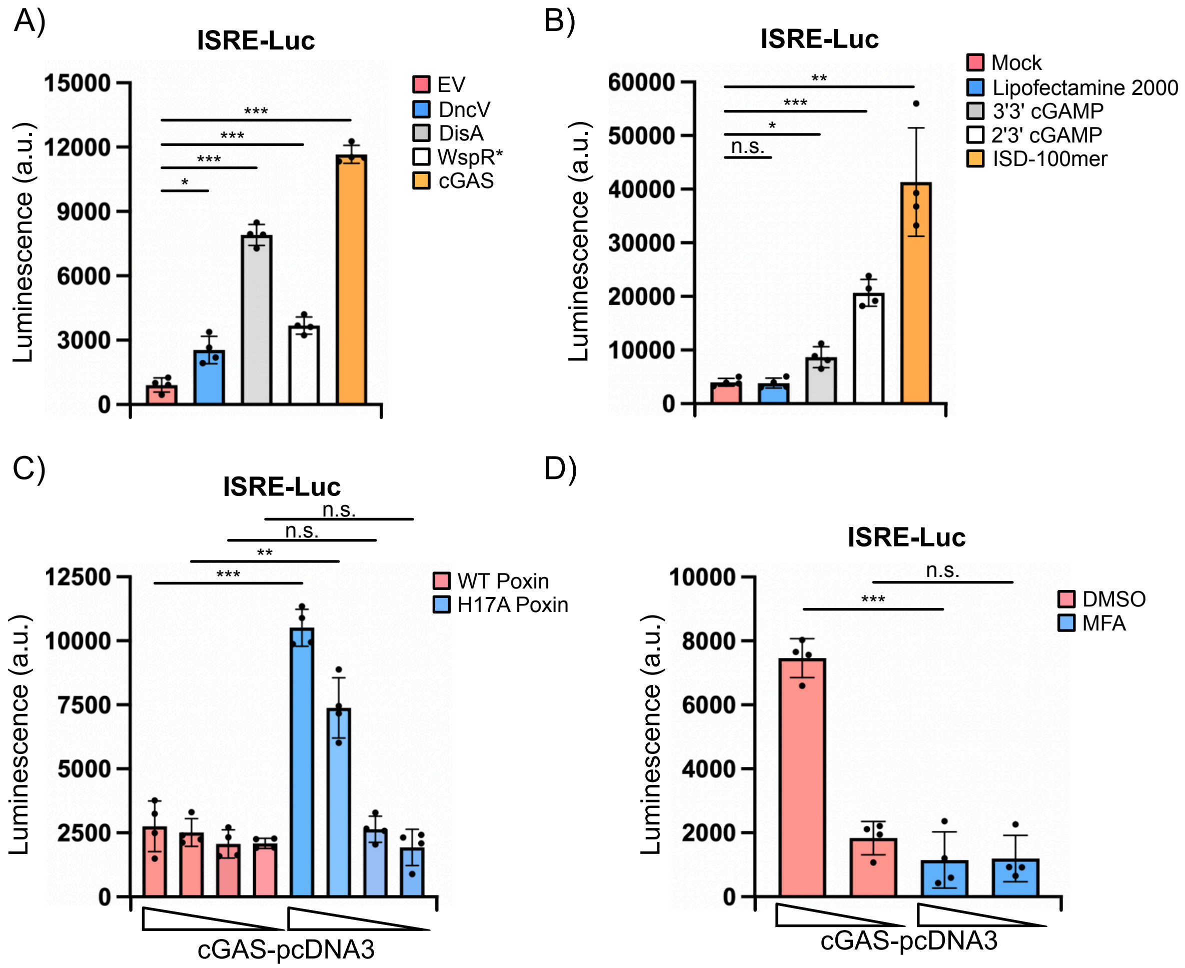

**Figure S2: Cyclic dinucleotides including 2'3' cGAMP are transferred from transfected cells into L929-ISRE-LUC reporter cells by co-culturing via gap junctions. (A)** Co-culture of L929-ISRE-LUC cells with HEK293T cells transfected with pcDNA3 empty vector (EV) or CDN cyclases. **(B)** L929-ISRE-LUC cells were mock treated, mock transfected with lipofectamine 2000, or transfected with 50 μM 3'3' cGAMP, 50 μM 2'3' cGAMP, or 1 μg ISD-100mer. Luciferase activity was quantified 18 h later. **(C)** Co-culture of L929-ISRE-LUC cells with HEK293T cells transfected with decreasing concentrations of cGAS-pcDNA3 (1000, 100, 10, 0 ng) and 3 μg of either wildtype (WT) or catalytically inactive (H17A) Poxin. **(D)** Co-culture of L929-ISRE-LUC cells with HEK293T cells transfected with cGAS-pcDNA3 (1000 or 0 ng) in the presence or absence of the gap junction inhibitor meclofenamic acid (MFA) (100 uM). $n = 4$ biological replicates.

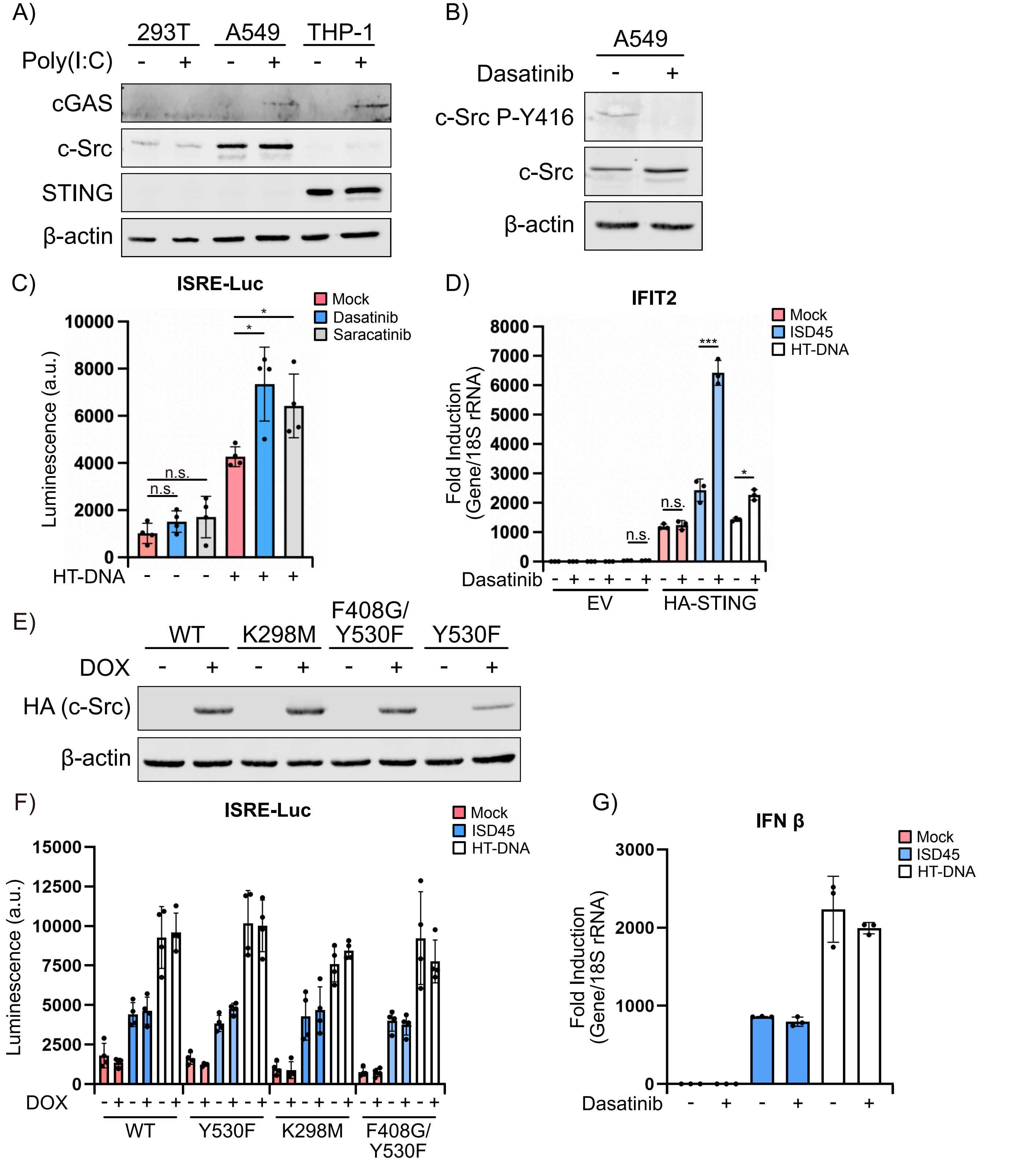

**Figure S3: c-Src, cGAS, and STING are expressed at different levels in various cell lines and impacts whether c-Src inhibition or overexpression regulates DNA sensing. (A)** Western blot of HEK293T, A549, and THP-1 cells treated with 1 μg/ml poly(I:C) for 16 h. b -actin was used as a loading control. **(B)** Western blot of A549 cells treated with 30 nM dasatinib or vehicle for 48 h. p Y416 is a marker for c-Src autophosphorylation and activation. b -actin was used as a loading control. **(C)** Co-culture of L929-ISRE-LUC cells with A549 cells treated with vehicle, dasatinib (30nM), or saracatinib (80 nM) for 48 h followed by transfection with HT-DNA for 6 h. **(D)** RT-qPCR analysis of IFIT2 levels from A549 cells transfected with EV or HA-STING pcDNA3 and treated with vehicle or dasatinib for 48 h followed by transfection with indicated DNA ligands for 6 h. All samples were normalized to 18S rRNA. **(E)** Western blot of A549 WT and mutant HA-c-Src-tet-RB SB cells treated with 1 μg/ml doxycycline (DOX) for 48 h. b -actin was used as a loading control. **(F)** Co-culture of L929-ISRE-LUC cells with cells in (E) followed by transfection with indicated DNA ligands for 6 h. **(G)** RT-qPCR analysis of IFN b levels from THP-1 cells treated with vehicle or dasatinib for 48 h followed by transfection with indicated DNA ligands for 6 h. All samples were normalized to 18S rRNA. *n*= 4 biological replicates for luciferase assay, and *n*= 3 biological replicates and representative qPCR with technical triplicates is shown.

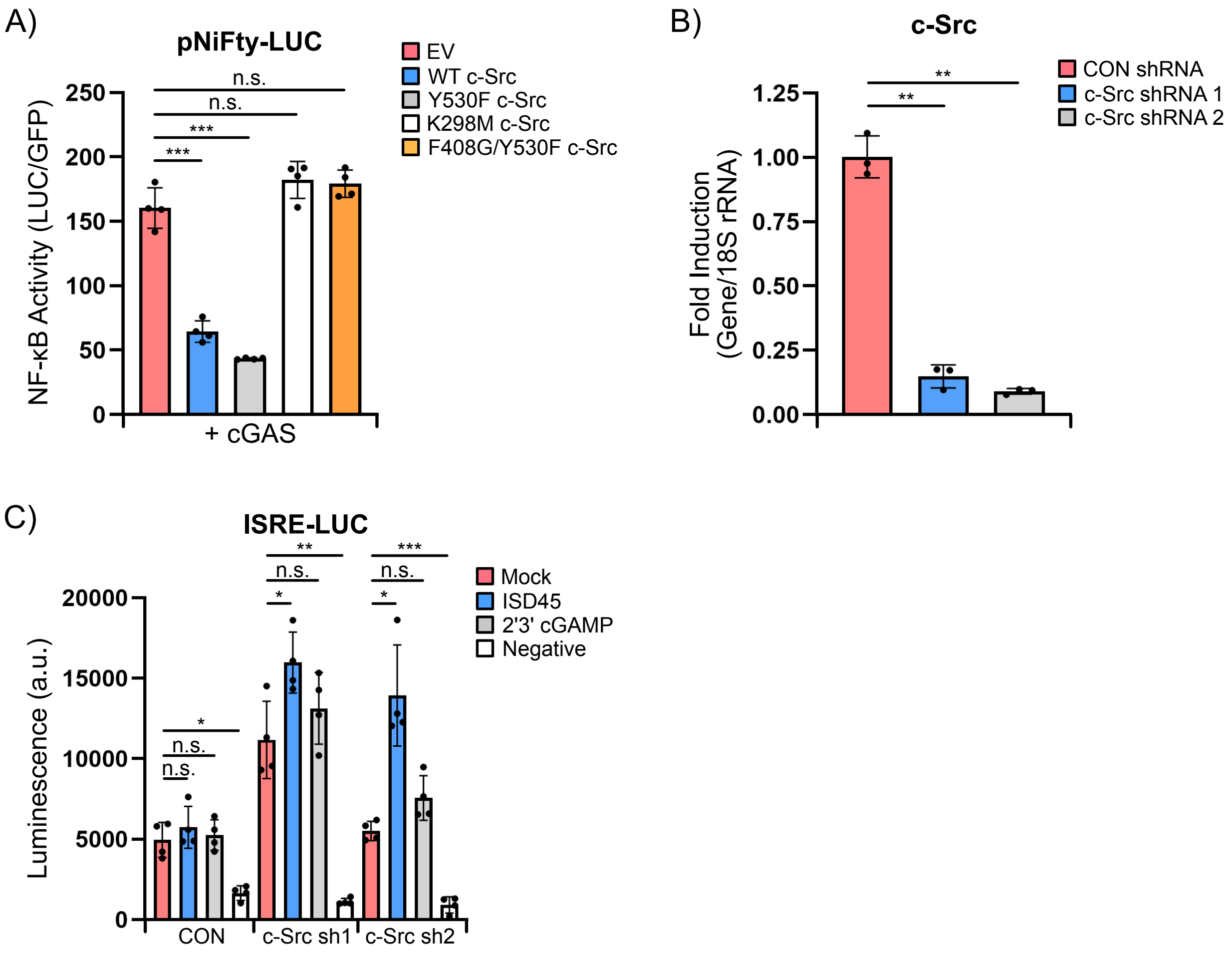

**Figure S4: c-Src inhibits cGAS-dependent NF-kB activity and c-Src depletion enhances a DNA-dependent IFN response. (A)** NF-kB luciferase assay from HEK293T cells transfected with pNiFty-LUC, cGAS pcDNA3, and EV or c-Src-HA pcDNA3 WT or mutant pcDNA3 for 24 h. NF-kB activity was analyzed by luciferase activity and normalized to transfected eGFP plasmid. **(B)** RT-qPCR analysis of c-Src expression in HEK293T cells transduced with DOX-inducible non-target (CON) or independent c-Src-targeting shRNAs. DOX was added for 72 h. c-Src levels in CON were set to 1. All samples were normalized to 18S rRNA. **(C)** Co-culture of L929-ISRE-LUC cells with cells in (B) transfected with cGAS pcDNA3 for 24 h followed by transfection of ISD45 or 2’3’ cGAMP for 6 h. Cells not transfected with cGAS were used as a negative control (Negative). *n*= 4 biological replicates for luciferase assays, and *n*= 3 biological replicates and representative qPCR with technical triplicates is shown.

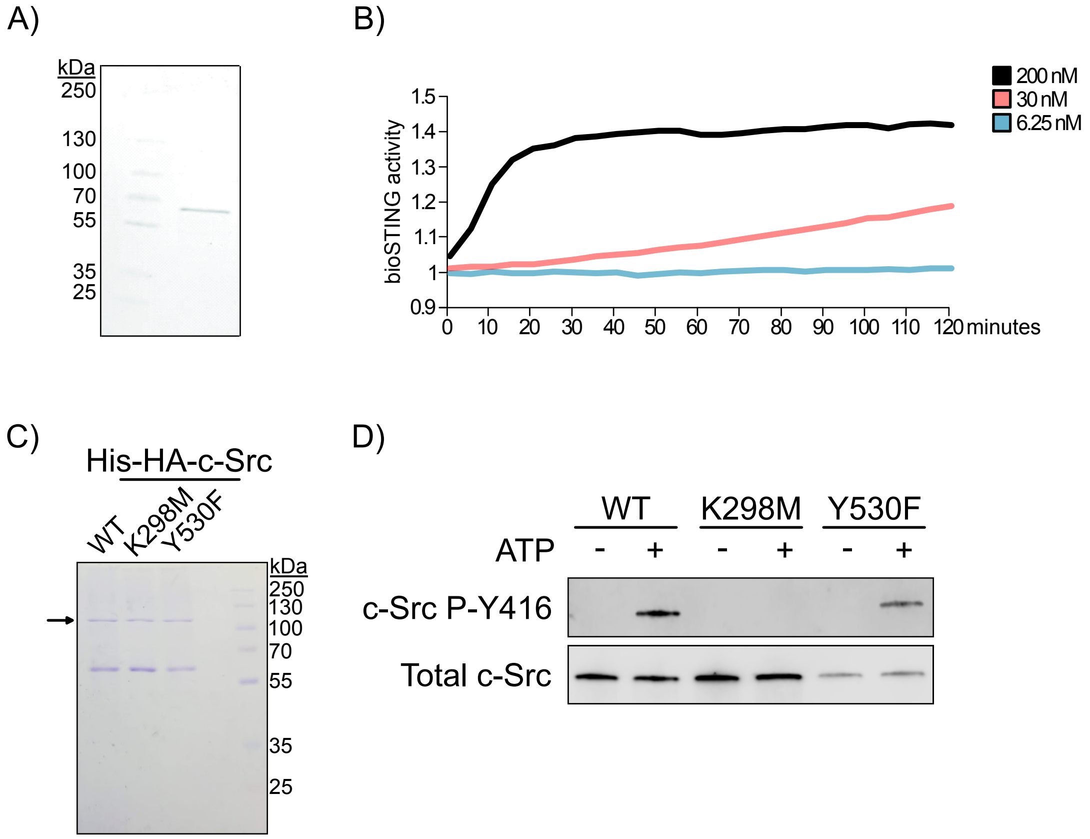

**Figure S5: Human FLAG-cGAS and HA-c-Src purification and activity validation. (A)** Coomassie brilliant blue stain of purified FLAG-cGAS after SEC and concentration. **(B)** BioSTING FRET assay using various concentration of purified FLAG-cGAS for 2 h. **(C)** Coomassie brilliant blue stain of purified WT and mutant HA-c-Src after concentration. Arrow depicts not-specific band. **(D)** Western blot of purified WT and mutant HA-c-Src *in vitro* kinase assays using cold ATP. c-Src Y416 designates autophosphorylation and c-Src kinase activity. *n*= 2 biological replicates.
