## Supplementary material for "The proto-oncogene c-Src phosphorylates cGAS to reduce DNA binding and activation": Table S1

| Oligos-qPCR | Sequence |
|---|---|
| 18S rRNA F | 5’ GTAACCCGTTGAACCCCATT |
| 18S rRNA R | 5’ CCATCCAATCGGTAGTAGCG |
| IFN Beta F | 5' CAGCAATTTTCAGTGTCAGAAGC |
| IFN Beta R | 5' TCATCCTGTCCTTGAGGCAGT |
| IFIT2 F | 5' AAATGCCATTTCACCTGGAACTTG |
| IFIT2 R | 5' GCTTTGAATTCACGATTCTGAAAC |
| TNF Alpha F | 5' CTCTTCTGCCTGCTGCACTTTG |
| TNF Alpha R | 5' ATGGGCTACAGGCTTGTCACTC |
| c-Src F | 5' CTGCTTTGGCGAGGTGTGGATG |
| c-Src R | 5' CCACAGCATACAACTGCACCAG |
| Murine IFN Beta F | 5' GCCTTTGCCATCCAAGAGATGC |
| Murine IFN Beta R | 5' ACACTGTCTGCTGGTGGAGTTC |
| Murine HPRT F | 5' CTGGTGAAAAGGACCTCTCGAAG |
| Murine HPRT R | 5' CCAGTTTCACTAATGACACAAACG |
| pc1 F | 5' GGAGATCTCCCGATCCCCTAT |
| pc1 R | 5' AGGGAGCAGATACTGGCTTAA |
| pc 2 F | 5' AAGACAATAGCAGGCATGCTG |
| pc2 R | 5' GCTACAGGGCGCGTGGGGATA |
| pc3 F | 5' CGGATCTGATCAAGAGACAGG |
| pc3 R | 5' AATAGCCTCTCCACCCAAGCG |
| GAPDH F | 5' GTCTCCTCTGACTTCAACAGCG |
| GAPDH R | 5' ACCACCCTGTTGCTGTAGCCAA |
| **Oligos-Cloning** | **Sequence** |
| cGAS Guide 1 lentiCRISPR V2 F | 5' CACCGGGCTTCCGCACGGAATGCCA |
| cGAS Guide 1 lentiCRISPR V2 R | 5' AAACTGGCATTCCGTGCGGAAGCCC |
| cGAS Guide 2 lentiCRISPR V2 F | 5' CACCGGGCCATGCAGAGAGCTTCCG |
| cGAS Guide 2 lentiCRISPR V2 R | 5' AAACCGGAAGCTCTCTGCATGGCCC |
| c-Src HA pcDNA3 F | 5' GGCCAAGCTTATGGGTAGCAACAAGAGCAAGCC |
| c-Src HA pcDNA3 R | 5' GGCCCTCGAGTCAAGCGTAATCTGGAACATCGTATGGGTACAAGAG GTTCTCCCCGGGC |
| c-Src Y530F SDM HA pcDNA3 F | 5' CCACCGAGCCCCAGTTCCAGCCCGGGGAGAAC |
| c-Src Y530F SDM HA pcDNA3 R | 5' GTTCTCCCCGGGCTGGAACTGGGGCTCGGTGG |
| c-Src K298M SDM HA pcDNA3 F | 5' GTACCACCAGGGTGGCCATCATGACCCTGAAGCCTGGCACGAT |
| c-Src K298M SDM HA pcDNA3 R | 5' ATCGTGCCAGGCTTCAGGGTCATGATGGCCACCCTGGTGGTAC |

| | |
|---|---|
| c-Src F408G SDM HA pcDNA3 F | 5' TGGTGTGCAAAGTGGCGGACGGTGGGCTGGCTCGGCTCATTGA |
| c-Src F408G SDM HA pcDNA3 R | 5' TCAATGAGCCGAGCCAGCCCACCGTCCGCCACTTTGCACACCA |
| HA-c-Src pSB-tet-RB F | 5' GCGGCCTCTGAGGCCATGTACCCATACGATGTTCCAGATTACGCTA TGGGTAGCAACAAGAGCAAGC |
| HA-c-Src pSB-tet-RB R | 5' GCGGCCTGACAGGCCCTAGAGGTTCTCCCCGGGCTG |
| His-HA-c-Src pSB-tet-RB F | 5' TCGAAAGGCCTCTGAGGCCATGCACCATCACCATCACCATATGTA CCCATACGATGTTCC |
| His-HA-c-Src pSB-tet-RB R | 5' GGAACATCGTATGGGTACATATGGTGATGGTGATGGTGCATGGCCTC AGAGGCCTTTCGA |
| FLAG-cGAS pcDNA3 F | 5' GCAAGCTTATGGACTACAAAGACGATGACGACAAGATGCAGCCTTGG CACGGAAAG |
| FLAG-cGAS pcDNA3 R | 5' GCCTCGAGTCAAAATTCATCAAAAACTGG |
| FLAG-cGAS Y248F SDM pcDNA3 F | 5' AATATTCCAACACTCGTGCATTCTACTTTGTGAAATTTAAAAG |
| FLAG-cGAS Y248F SDM pcDNA3 R | 5' CTTTTAAATTTCACAAAGTAGAATGCACGAGTGTTGGAATATT |
| FLAG-cGAS Y248E SDM pcDNA3 F | 5' AATATTCCAACACTCGTGCAGAGTACTTTGTGAAATTTAAAAG |
| FLAG-cGAS Y248E SDM pcDNA3 R | 5' CTTTTAAATTTCACAAAGTACTCTGCACGAGTGTTGGAATATT |
| c-Src g1 pLKO-tet-ON F | 5' CCGGCATCCTCAGGAACCAACAATTCTCGAGAATTGTTGGTTCCTG AGGATGTTTTTG |
| c-Src g1 pLKO-tet-ON R | 5' AATTCAAAAACATCCTCAGGAACCAACAATTCTCGAGAATTGTTGGT TCCTGAGGATG |
| c-Src g2 pLKO-tet-ON F | 5' CCGGCTGACTGAGCTCACCACAAAGCTCGAGCTTTGTGGTGAGCT CAGTCAGTTTTTG |
| c-Src g2 pLKO-tet-ON R | 5' AATTCAAAAACTGACTGAGCTCACCACAAAGCTCGAGCTTTGTGGT GAGCTCAGTCAG |
| Oligos-TOPO PCR | Sequence |
| cGAS g1 g2 PCR F | 5' TGAGCTTCAACTTCTCCAAAACCG |
| cGAS g1 g2 PCR F | 5' CGAGACTTTTGTAGCCTCAGGAAAG |
| Antibodies | Company; Dilution |
| HA | Abcam 18181 |
| STING | CST 13647S |
| Beta actin | CST 3700S |
| cGAS | CST 15102S |
| FLAG | Millipore F1804 |
| c-Src | CST 2109S |
| c-Src phospho Y416 | CST 2101S |
| Plasmids | Source |
| pJP1520-SRC | DNASU |
| HA-STING pcDNA3 | Dr. Genhong Cheng |
| cGAS pcDNA3 | Dr. Genhong Cheng |

| | |
|---|---|
| pSB-tet-RB | Addgene |
| SB100X | Addgene |
| lentiCRISPR V2 | Addgene |
| psPAX2 | Addgene |
| pVSV-G | Addgene |
| pSUMO2 human cGAS-FL | Dr. Philip Kranzusch |
| pET15b-bioSTING | Pollock et al. PMID: 32669552 |
| DncV pcDNA3 | Dr. Philip Kranzusch |
| DisA pcDNA3 | Dr. Philip Kranzusch |
| WspR* pcDNA3 | Dr. Philip Kranzusch |
| WT Poxin pcDNA3 | Dr. Philip Kranzusch |
| Poxin H17A pcDNA3 | Dr. Philip Kranzusch |
| pNiFty-LUC | Invivogen |
| eGFP pcDNA3 | Addgene |
| pLKO-tet-ON | Addgene |
| HA-c-Src WT pSB-tet-RB | This paper |
| HA-c-Src Y530F pSB-tet-RB | This paper |
| HA-c-Src F408G/Y530F pSB-tet-RB | This paper |
| HA-c-Src K298M pSB-tet-RB | This paper |
| c-Src WT HA pcDNA3 | This paper |
| c-Src Y530F HA pcDNA3 | This paper |
| c-Src F408G/Y530F HA pcDNA3 | This paper |
| c-Src K298M HA pcDNA3 | This paper |
| FLAG-cGAS pcDNA3 | This paper |
| Scramble lentiCRISPR V2 | Dunker et al. PMID: 33852834 |
| cGAS Guide 1 lentiCRISPR V2 | This paper |
| cGAS Guide 2 lentiCRISPR V2 | This paper |
| pSUMO2 human FLAG-cGAS-FL | This paper |
| His-HA-c-Src WT pSB-tet-RB | This paper |
| His-HA-c-Src Y530F pSB-tet-RB | This paper |
| His-HA-c-Src K298M pSB-tet-RB | This paper |
| FLAG-cGAS Y248F SDM pcDNA3 | This paper |

| | |
|---|---|
| FLAG-cGAS Y248E SDM pcDNA3 | This paper |
| c-Src g1 pLKO-tet-ON | This paper |
| c-Src g2 pLKO-tet-ON | This paper |
